# SMNN: Batch Effect Correction for Single-cell RNA-seq data via Supervised Mutual Nearest Neighbor Detection

**DOI:** 10.1101/672261

**Authors:** Yuchen Yang, Gang Li, Huijun Qian, Kirk C. Wilhelmsen, Yin Shen, Yun Li

**Affiliations:** Department of Genetics, University of North Carolina, Chapel Hill, NC 27599, USA; Statistics and Operations Research, University of North Carolina, Chapel Hill, NC 27599, USA; Biostatistics, University of North Carolina, Chapel Hill, NC 27599, USA; Computer Science, University of North Carolina, Chapel Hill, NC 27599, USA; Renaissance Computing Institute, University of North Carolina, Chapel Hill, NC 27599, USA; Institute for Human Genetics, University of California, San Francisco, San Francisco, CA 94143, USA; Department of Neurology, University of California, San Francisco, San Francisco, CA 94143, USA

**Keywords:** single-cell RNA sequencing, batch effect, supervised mutual nearest neighbor

## Abstract

Batch effect correction has been recognized to be indispensable when integrating single-cell RNA sequencing (scRNA-seq) data from multiple batches. State-of-the-art methods ignore single-cell cluster label information, but such information can improve effectiveness of batch effect correction, particularly under realistic scenarios where biological differences are not orthogonal to batch effects. To address this issue, we propose SMNN for batch effect correction of scRNA-seq data via supervised mutual nearest neighbor detection. Our extensive evaluations in simulated and real datasets show that SMNN provides improved merging within the corresponding cell types across batches, leading to reduced differentiation across batches over MNN, Seurat v3, and LIGER. Furthermore, SMNN retains more cell type-specific features, partially manifested by differentially expressed genes identified between cell types after SMNN correction being biologically more relevant, with precision improving by up to 841%.

**Key Points:**

1. Batch effect correction has been recognized to be critical when integrating scRNA-seq data from multiple batches due to systematic differences in time points, generating laboratory and/or handling technician(s), experimental protocol, and/or sequencing platform.
2. Existing batch effect correction methods that leverages information from mutual nearest neighbors across batches (for example, implemented in SC3 or Seurat) ignore cell type information and suffer from potentially mismatching single cells from different cell types across batches, which would lead to undesired correction results, especially under the scenario where variation from batch effects is non-negligible compared with biological effects.
3. To address this critical issue, here we present SMNN, a supervised machine learning method that first takes cluster/cell-type label information from users or inferred from scRNA-seq clustering, and then searches mutual nearest neighbors within each cell type instead of global searching.
4. Our SMNN method shows clear advantages over three state-of-the-art batch effect correction methods and can better mix cells of the same cell type across batches and more effectively recover cell-type specific features, in both simulations and real datasets.

## Introduction

An ever-increasing amount of single cell RNA-sequencing (scRNA-seq) data has been generated as scRNA-seq technologies mature and sequencing costs continue dropping. However, large scale scRNA-seq data, for example, those profiling tens of thousands to millions of cells (such as the Human Cell Atlas Project [1], almost inevitably involve multiple batches across time points, laboratories, or experimental protocols. The presence of batch effect renders joint analysis across batches challenging [2, 3]. Batch effect, or systematic differences in gene expression profiles across batches, not only can obscure the true underlying biology, but also may lead to spurious findings. Thus, batch effect correction, which aims to mitigate the discrepancies across batches, is crucial and deemed indispensable for the analysis of scRNA-seq data across batches [4].

Because of its importance, a number of batch effects correction methods has been recently proposed and implemented. Most of these methods, including limma [5], ComBat [6], and svaseq [7], are regression-based. Among them, limma and ComBat explicitly model known batch effect as a blocking term. Because of the regression framework adopted, standard statistical approaches to estimate the regression coefficients corresponding to the blocking term can be conveniently employed. In contrast, svaseq is often used to detect underlying unknown factors of variation, for instance, unrecorded differences in the experimental protocols. svaseq first identifies these unknown factors as surrogate variables and subsequently corrects them. For these regression-based methods, once the regression coefficients are estimated or the unknown factors are identified, one can then regress out these batch effects accordingly, obtaining residuals that will serve as the batch-effect corrected expression matrix for further analyses. These methods have become standard practice in the analysis of bulk RNA-seq data. However, when it comes to scRNA-seq data, one key underlying assumption behind these methods, that the cell composition within each batch is identical, might not hold. Consequently, estimates of the coefficients might be inaccurate. As a matter of fact, when applied to scRNA-seq data, the corrected results derived from these methods widely adopted for bulk RNA-seq data might be even inferior to raw data without no correction, in some extreme cases [8].

To address the heterogeneity and high dimensionality of complex data, several dimension-reduction approaches have been adopted. An incomplete list of these strategies includes principal component analysis (PCA), autoencoder, or force-based methods such as t-distributed stochastic neighbor embedding (t-SNE) [9]. Through those dimension reduction techniques, one can project new data onto the reference dataset using a set of landmarks from the reference [8, 10-12] to remove batch effects between any new dataset and the reference dataset. Such projection methods require the reference batch contains all the cell types across batches. As one example, Spitzer *et al*. [11] employed force-based dimension reduction and showed that leveraging a few landmark cell types from bone marrow (the most appropriate tissue in that it provides the most complete coverage of immune cell types) allowed mapping and comparing immune cells across different tissues and species. When applied to scRNA-seq data, however, these methods suffer when cells from a new batch fall out of the space inferred from the reference. Furthermore, determining the dimensionality of the low dimensional manifolds is still an open and challenging problem. To address the limitations of existing methods, two recently developed batch effect correction methods, MNN and Seurat v3, adopt the concept of leveraging information of mutual nearest neighbors (MNN) across batches [8, 12], and demonstrate superior performance over alternative methods [8, 12]. However, this MNN-based strategy ignores cell type information and suffers from potentially mismatching cells from different cell types/states across batches, which may lead to undesired correction results. For example, under the scenario depicted in **Fig. 1b**, MNN leads to cluster 1 (C1) and cluster 2 (C2) mis-corrected due to mismatching single cells in the two clusters/cell-types across batches.

**Fig. 1.**
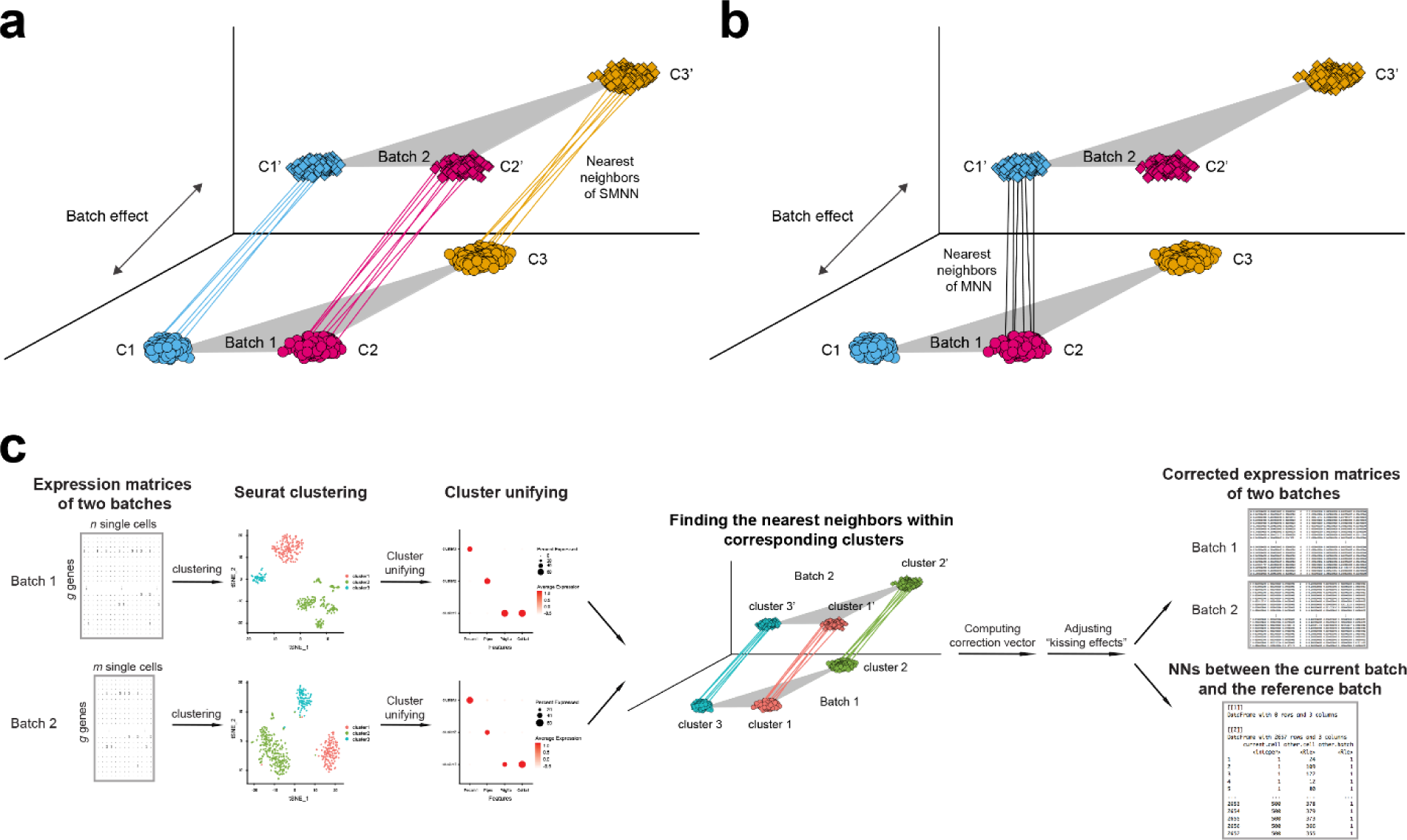
Overview of SMNN. Schematics for detecting mutual nearest neighbors between two batches under a non-orthogonal scenario **(a)** in SMNN; and **(b) in** MNN. **(c)** Workflow of SMNN. Single cell clustering is first performed within each batch using Seurat v3; and then SMNN takes user-specified marker gene information for each cell type to match clusters/cell types across batches. With the clustering and cluster-specific marker gene information, SMNN searches mutual nearest neighbors within each cell type and performs batch effect correction accordingly.

To address the above issue, here we present SMNN, a supervised machine learning method that explicitly incorporates cell type information. SMNN performs nearest neighbor searching within the same cell type, instead of global searching ignoring cell type labels (**Fig. 1a**). Cell type information, when unknown *a priori*, can be inferred via clustering methods [13-16].

## Results

### SMNN Framework

The motivation behind our SMNN is that single-cell cluster or cell type information has the potential aid the identification of most relevant nearest neighbors and subsequently improve batch effect correction. A preliminary clustering before any correction can provide knowledge regarding cell composition within each batch, which serves as the cellular correspondence across batches (**Fig. 1a**). With this clustering information, we can refine the nearest neighbor searching space within a certain population of cells that are of the same or similar cell type(s) or state(s) across all batches.

SMNN takes a natural two-step approach to leverage cell type label information for enhanced batch effect correction (**Fig. 1c and Supplementary Section 1**). First, it takes the expression matrices across multiple batches as input, and performs clustering separately for each batch. Specifically, in this first step, SMNN uses Seurat v3 [17] where dimension reduction is conducted via principal component analysis (PCA) to the default of 20 PCs, and then graph-based clustering follows on the dimension-reduced data with *resolution* parameter of 0.9 [18, 19]. Obtaining an accurate matching of the cluster labels across batches is of paramount importance for subsequent nearest neighbor detection. SMNN requires users to specify a list of marker genes and their corresponding cell type labels to match clusters/cell types across batches. We hereafter refer to this cell type or cluster matching as cluster harmonization across batches. Because not all cell types are necessarily shared across batches, and no prior knowledge exists regarding the exact composition of cell types in each batch, SMNN allows users to take discretion in terms of the marker genes to include, representing the cell types that are believed to be shared across batches. Based on the marker gene information, a harmonized label is assigned to *every* cluster identified across all the batches according to two criteria: the percentage of cells in a cluster expressing a certain marker gene and the average gene expression levels across all the cells in the cluster. After harmonization, cluster labels are unified across batches. This completes step one of SMNN. Note that if users have *a priori* knowledge regarding the cluster/cell-type labels, the clustering step could be bypassed completely.

With the harmonized cluster or cell type label information obtained in the first step, SMNN, in the second step, searches mutual nearest neighbors only within each matched cell type between the first batch (which serves as the reference batch) and any of the other batches (the current batch), and performs batch effect correction accordingly. Compared to MNN or Seurat v3, where the mutual nearest neighbors or anchor cells are searched globally, SMNN identifies neighbors from the same cell population or state. After mutual nearest neighbors are identified, similar to MNN, SMNN first computes batch effect correction vector for each identified pair of cells, and then calculates, for each cell, the cell-specific correction vectors by exploiting a Gaussian kernel to obtain a weighted average across all the pair-specific vectors with mutual nearest neighbors of the cell under consideration. The correction vectors obtained from shared cell-types will be applied to correct all cells including those belonging to batch-specific cell types (detailed in **Supplementary Section 2**) Each cell’s correction vector is further scaled according to the cell’s location in the space defined by the correction vector, and standardized according to quantiles across batches, in order to eliminate “kissing effects”. “Kissing effects” refer to the phenomenon that only the surfaces of cell-clouds across batches are brought in contact (rather than fully merged), commonly observed with naïve batch effect correction [8] (an example detailed in **Supplementary Section 3** and visualized in **Supplementary Fig. S1**). At the end of the second step, SMNN returns the batch-effect corrected expression matrix including all genes from the input matrix for each batch, as well as the information regarding nearest neighbors between the reference batch and the current batch under correction. This step is carried out for every batch other than the reference batch so that all batches are corrected to the same reference batch in the end.

### Simulation results

Since MNN has been shown to excel alternative methods [4, 8], we here focus on comparing our SMNN with MNN. We first compared the performance of SMNN to MNN in simulated data. In our simulations, SMNN demonstrates superior performance over MNN under both orthogonal and non-orthogonal scenarios (**Fig. 2 and 3 and Supplementary Fig. S2-4**). We show t-SNE plot for each cell type before and after MNN and SMNN correction under both the orthogonal and non-orthogonal scenarios. Under orthogonality, the two batches partially overlapped in the t-SNE plot before correction, suggesting that the variation due to batch effect was indeed much smaller than that due to biological effect. Both MNN and SMNN successfully mixed single cells from two batches (**Supplementary Fig. S3**). However, for cell types 1 and 3, there were still some cells from the second batch left unmixed with those from the first batch after MNN correction (**Supplementary Fig. S3a and c**). Under the non-orthogonal scenario, the differences between two batches were more pronounced before correction, and SMNN apparently outperformed MNN (**Supplementary Fig. S4**), especially in cell type 1 (**Supplementary Fig. S4a**). Moreover, we also computed Frobenius norm distance [20] for each cell between its simulated true profile before introducing batch effects and after SMNN and MNN correction. The results showed an apparently reduced deviation from the truth after SMNN correction than MNN (**Fig. 3**). We have also simulated data using the original simulation framework in Haghverdi *et al*. [8], which does not allow precise control of orthogonality (detailed in **Method Section**) and seems to simulate data closer to those under orthogonal cases (**Supplementary Fig. S5a**). Applying SMNN and MNN to such simualted data, we also found that SMNN showed slight advantages (**Supplementary Fig. S5b**) These results suggest that SMNN provides improved batch effect correction over MNN under both orthogonal and non-orthogonal scenarios.

**Fig. 2.**
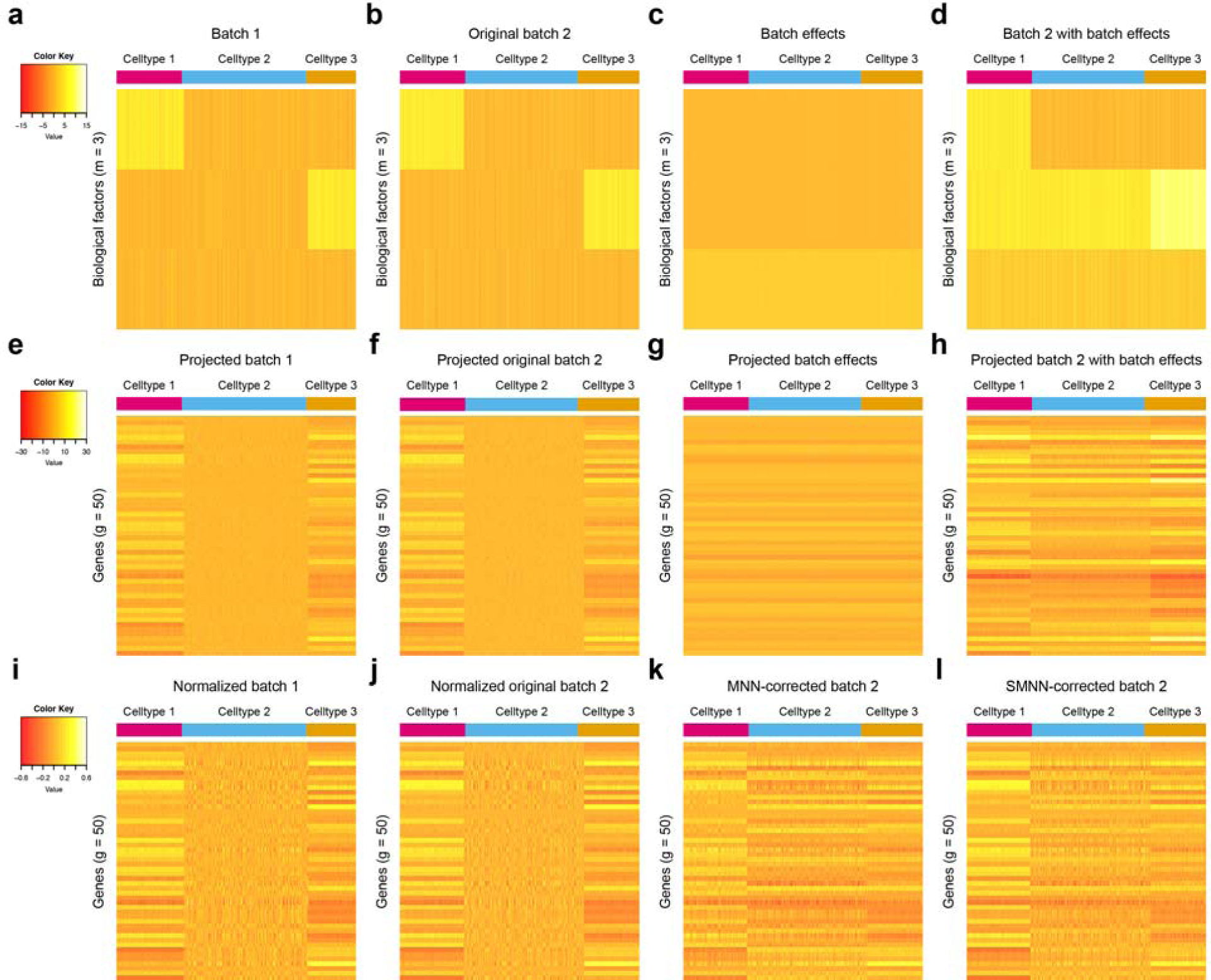
Heatmap of gene expression matrices for simulated data under non-orthogonal scenario. **(a), (b), (c)** and **(d)** show the 3-dimensional biological space with rows of each heatmap representing biological factors and columns corresponding to single cells. **(e), (f), (g)** and **(h)** show the high dimensional gene expression profiles with rows corresponding to genes and columns again representing single cells. **(a), (e)** and **(i)** correspond to the batch 1, and **(b), (f)** and **(j)** correspond to batch 2. **(c)** and **(g)** provide a visualization for the direction of batch effects in low-dimension biological space and high-dimension gene expression spaces, respectively. **(d)** and **(h)**, sum of **(b)** and **(c)** and sum of **(f)** and **(g)** respectively, are “observed” data for cells in batch 2 in low and high dimensional space respectively. **(i)** and **(j)** are the cosine-normalized data for batch 1 and original batch 2. Note “original” is in the sense that no batch effects have been introduced to the data yet. **(k)** and **(l)** are the MNN and SMNN corrected results, respectively.

**Fig. 3.**
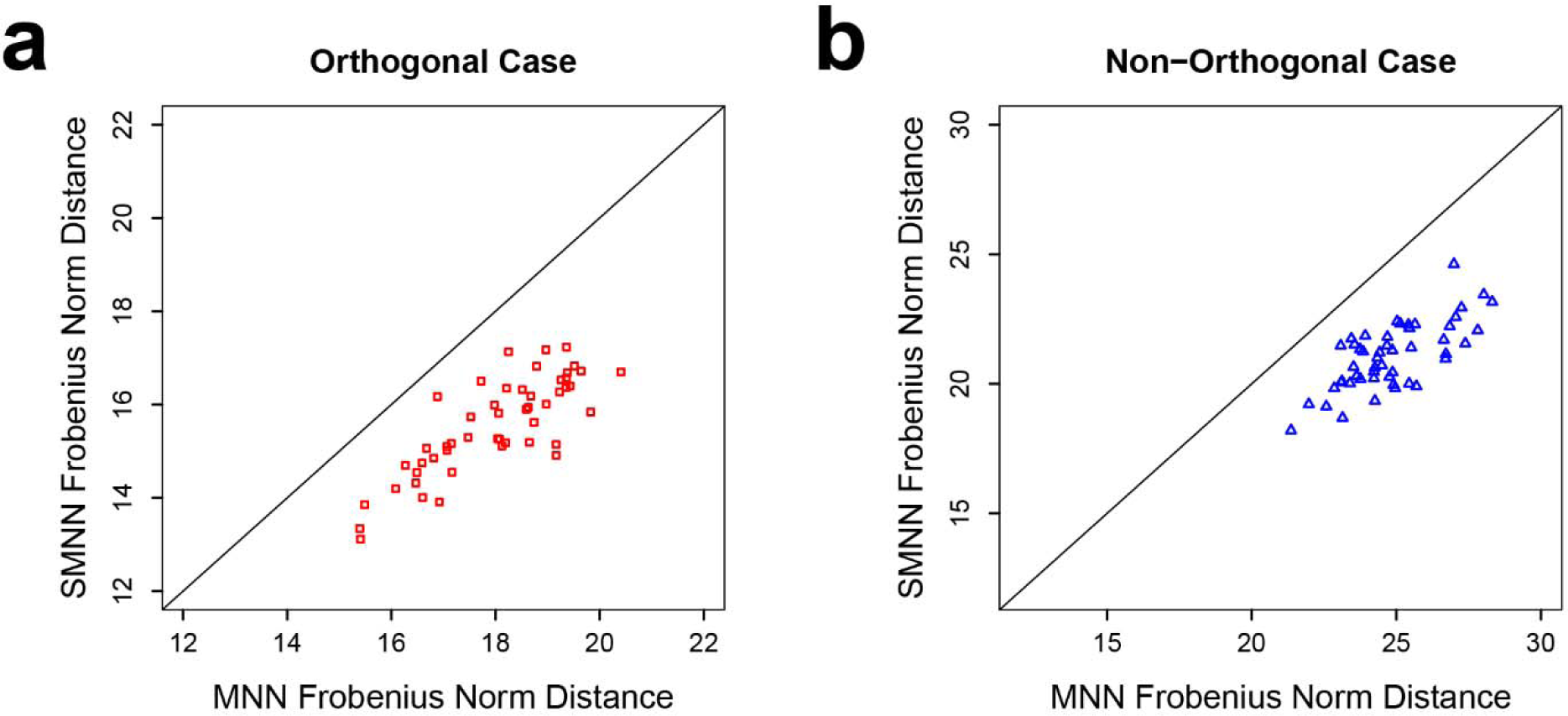
Frobenius norm distance between two batches after SMNN and MNN correction in simulation data under orthogonal (left) and non-orthogonal scenarios (right).

### Real data results

For performance evaluation in real data, we first carried out batch effect correction on two hematopoietic datasets (**Supplementary Table S1**) using four methods: our SMNN, published MNN, Seurat v3 and LIGER. **Fig. 4a-e** shows UMAP plot before and after correction. Notably, all four methods can substantially mitigate discrepancy between the two datasets. Comparatively, SMNN better mixed cells of the same cell type across batches than the other three methods, and seemed to better position cells from batch-specific cell types with respect to other biologically related cell types (**Supplementary Fig. S6 and S7**), especially for common myeloid progenitor (CMP) and megakaryocyte-erythrocyte progenitor (MEP) cells, which were wrongly corrected by MNN due to sub-optimal nearest neighbor search ignoring cell type information (**Supplementary Fig. S8**). Correspondingly, SMNN corrected data exhibits the lowest F value than that from the other three methods. Specifically, F value is with reduced by 81.5 - 96.6% on top of MNN, Seurat v3, and LIGER, respectively (**Fig. 4f**). Furthermore, we compared the distance for the cells between batch 1 and 2, and found that, compared to data before correction, both MNN and SMNN reduced the Euclidean distance between the two batches (**Supplementary Fig. S9**). In addition, SMNN further decreased the distance by up to 8.2% than MNN (2.8%, 4.3% and 8.2% for cells of type CMP, MEP and granulocyte-monocyte progenitor (GMP) cells, respectively). Under scenarios where we only have partial cell type information, SMNN still better mixed cells of the same cell type across batches (detailed in **Supplementary Section 3; Supplementary Fig. S10a-c and e-g**), and manifested the best/lowest F values, compared with uncorrected and MNN-corrected data (**Supplementary Fig. S10d and h**). These results suggest improved batch effect correction by SMNN, compared to unsupervised correction methods.

**Fig. 4.**
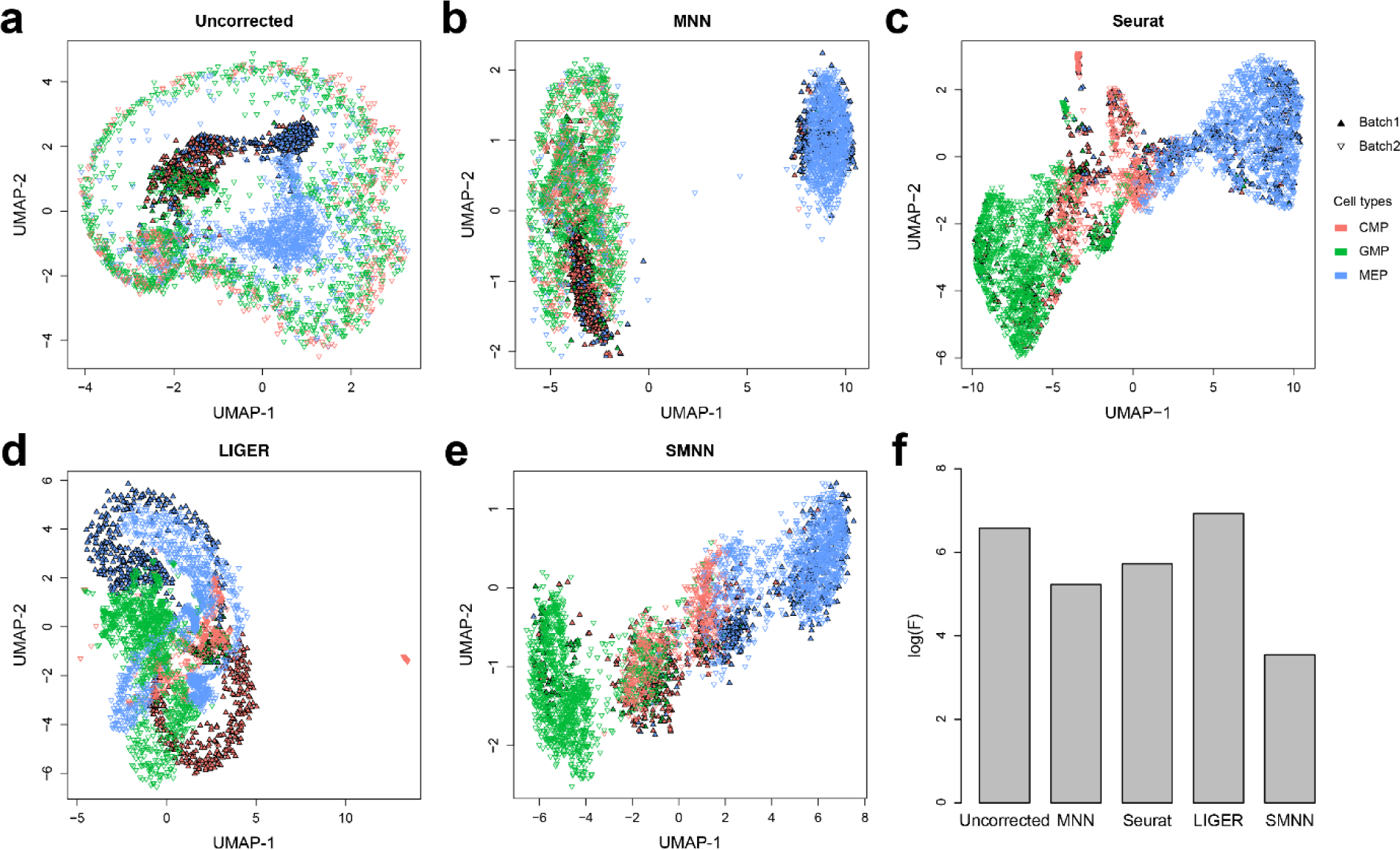
Performance comparison between SMNN and MNN in two hematopoietic datasets. (**a**) UMAP plots for two hematopoietic datasets before batch effect correction. Solid and inverted triangle represent the first and second batch, respectively; and different cell types are shown in different colors. (**b-e**) UMAP plots for the two hematopoietic datasets after correction with MNN, Seurat v3, LIGER, and SMNN. (**f**) Logarithms of F-statistics for merged data of the two batches.

### SMNN identifies differentially expressed genes that are biologically relevant

We then compared the differentially expressed genes (DEGs) among different cell types identified by SMNN and MNN. After correction, in the merged hematopoietic dataset, 1012 and 1145 up-regulated DEGs were identified in CMP cells by SMNN and MNN, respectively, when compared to GMP cells, while 1126 and 1108 down-regulated DEGs were identified by the two methods, respectively (**Fig. 5a and Supplementary Fig S11a**). Of them, 736 up-regulated and 842 down-regulated DEGs were shared between SMNN and MNN corrected data. Gene ontology (GO) enrichment analysis showed that, the DEGs detected only by SMNN were overrepresented in GO terms related to blood coagulation and hemostasis, such as platelet activation and aggregation, hemostasis, coagulation and regulation of wound healing (**Fig. 5b**). Similar DEG detection was carried out to detect genes differentially expressed between CMP and MEP cells. 181 SMNN-specific DEGs were identified out of the 594 up-regulated DEGs in CMP cells when compared to MEP cells (**Fig. 5c**), and they were found to be enriched for GO terms involved in immune cell proliferation and differentiation, including regulation of leukocyte proliferation, differentiation and migration, myeloid cell differentiation and mononuclear cell proliferation (**Fig. 5d**). Lastly, genes identified by SMNN to be up-regulated in GMP when compared to MEP cells, were found to be involved in immune processes; whereas up-regulated genes in MEP over GMP were enriched in blood coagulation (**Supplementary Fig. S11e-h**). Comparatively, the GO terms enriched for MNN-specific DEGs seem not particularly relevant to corresponding cell functions (**Supplementary Fig. S12**). These cell-function-relevant SMNN-specific DEGs indicate SMNN can maintain some cell features that are missed by MNN after correction.

**Fig. 5.**
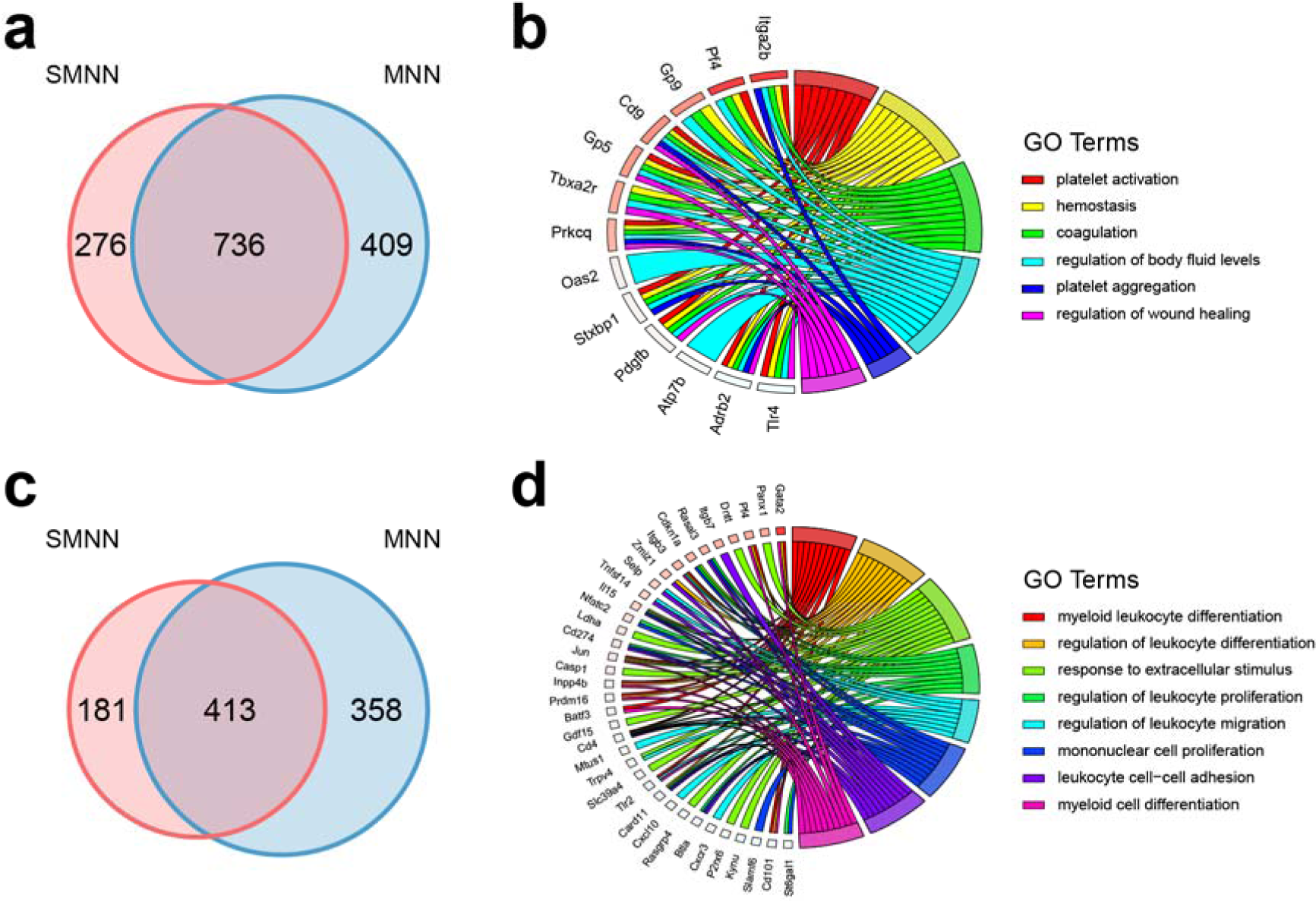
Comparison of differentially expressed genes (DEGs), identified in the merged dataset by pooling batch 1 data with batch 2 data after SMNN and MNN correction. (**a**) Overlap of DEGs up-regulated in CMP over GMP after SMNN and MNN correction. (**b**) Feature enriched GO terms and the corresponding DEGs up-regulated in CMP over GMP. (**c**) Overlap of DEGs up-regulated in CMP over MEP after SMNN and MNN correction. (**d**) Feature enriched GO terms and the corresponding DEGs up-regulated in CMP over MEP.

In addition, we considered two sets of “working truth”: first, DEGs identified in uncorrected batch 1; second DEGs identified in batch 2, and we compared SMNN and MNN results to both sets of working truth. The results showed that, in both comparisons (one comparison for each set of working truth), fewer DEGs were observed in SMNN-corrected batch 2, but higher precision and lower false negative rate in each of the three cell types than those in MNN results (**Fig. 6 and Supplementary Fig. S13-15**). When compared to the uncorrected batch 1, 3.6% - 841% improvements in precision were observed in SMNN results than MNN (**Fig. 6 and Supplementary Fig. S14**). Similarly, SMNN increased the precision by 6.2% - 54.0% on top of MNN when compared to uncorrected batch 2 (**Supplementary Fig. S15**). We also performed DEG analysis at various adjusted *p*-value thresholds and the results showed that the better performance of SMNN is not sensitive to the p-value cutoff we used for DEG detection (detailed in **Supplementary Section 3; Supplementary Fig. S16**). Such an improvement in the accuracy of DEG identification indicates that higher amount of information regarding cell structure was retained after SMNN correction than MNN.

**Fig. 6.**
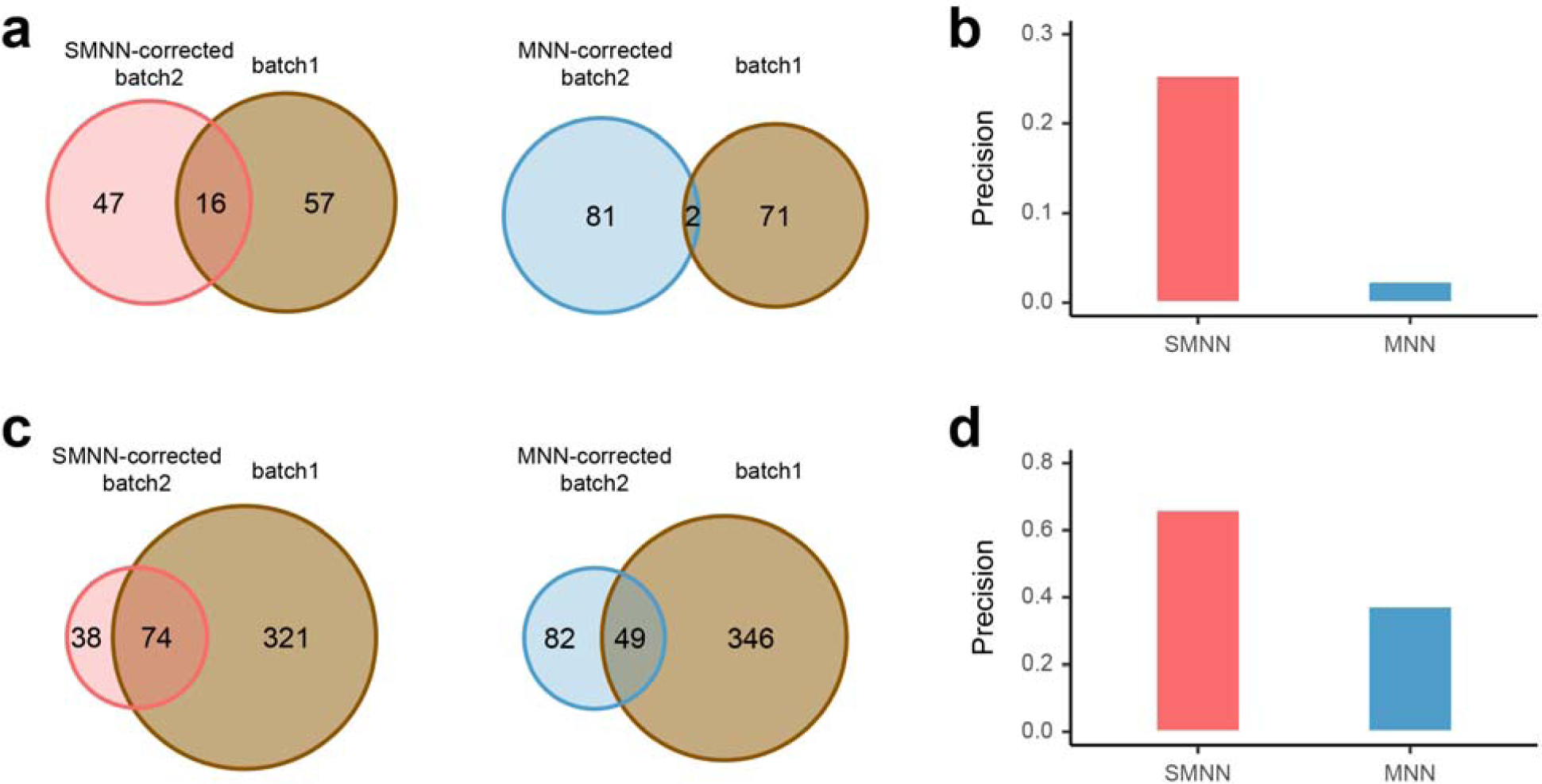
Reproducibility of DEGs (between CMP and GMP), identified in uncorrected batch 1 and in SMNN or MNN-corrected batch 2. (**a**) Reproducibility of DEGs up-regulated in CMP over GMP, detected in batch 1, versus SMNN (left) or MNN-corrected (right) batch 2. (**b**) True positive rate (TPR) of the DEGs (between CMP and GMP) identified in batch 2 after SMNN and MNN correction. (**c**) Reproducibility of DEGs up-regulated in GMP over CMP, identified in the uncorrected batch 1, and in SMNN (left) or MNN-corrected (right) batch 2. (**d**) TPR of the DEGs up-regulated in GMP over CMP identified in batch 2 after SMNN and MNN correction.

We also identified DEGs between T cells and B cells in the merged human peripheral blood mononuclear cells (PBMCs) and T cell datasets after SMNN and MNN correction, respectively (**Supplementary Fig. S17**). Compared to B cells, 3213 and 4180 up-regulated DEGs were identified in T cells by SMNN and MNN, respectively, 2203 of which were shared between the two methods (**Supplementary Fig. S17e**). GO enrichment analysis showed that, the SMNN-specific DEGs were significantly enriched for GO terms relevant to the processes of immune signal recognition and T cell activation, such as T cell receptor signaling pathway, innate immune response-activating signal transduction, cytoplasmic pattern recognition receptor signaling pathway and regulation of autophagy (**Supplementary Fig. S17f**). In B cells, 5422 and 3462 were found to be up-regulated after SMNN and MNN correction, where 2765 were SMNN-specific (**Supplementary Fig. S17g**). These genes were overrepresented in GO terms involved in protein synthesis and transport, including translational elongation and termination, ER to Golgi vesicle-mediated transport, vesicle organization and Golgi vesicle budding (**Supplementary Fig. S17h**). These results again suggest that SMNN more accurately retains or rescues cell features after correction.

### SMNN more accurately identifies cell clusters

Finally, we examined the ability to differentiate cell types after SMNN and MNN correction in three datasets (**Supplementary Table S1**). In all three real datasets, Adjusted Rand Index (ARI) after SMNN correction showed - 42.3% improvements over that of MNN (**Fig. 7**), suggesting that SMNN correction more effectively recovers cell-type specific features.

**Fig. 7.**
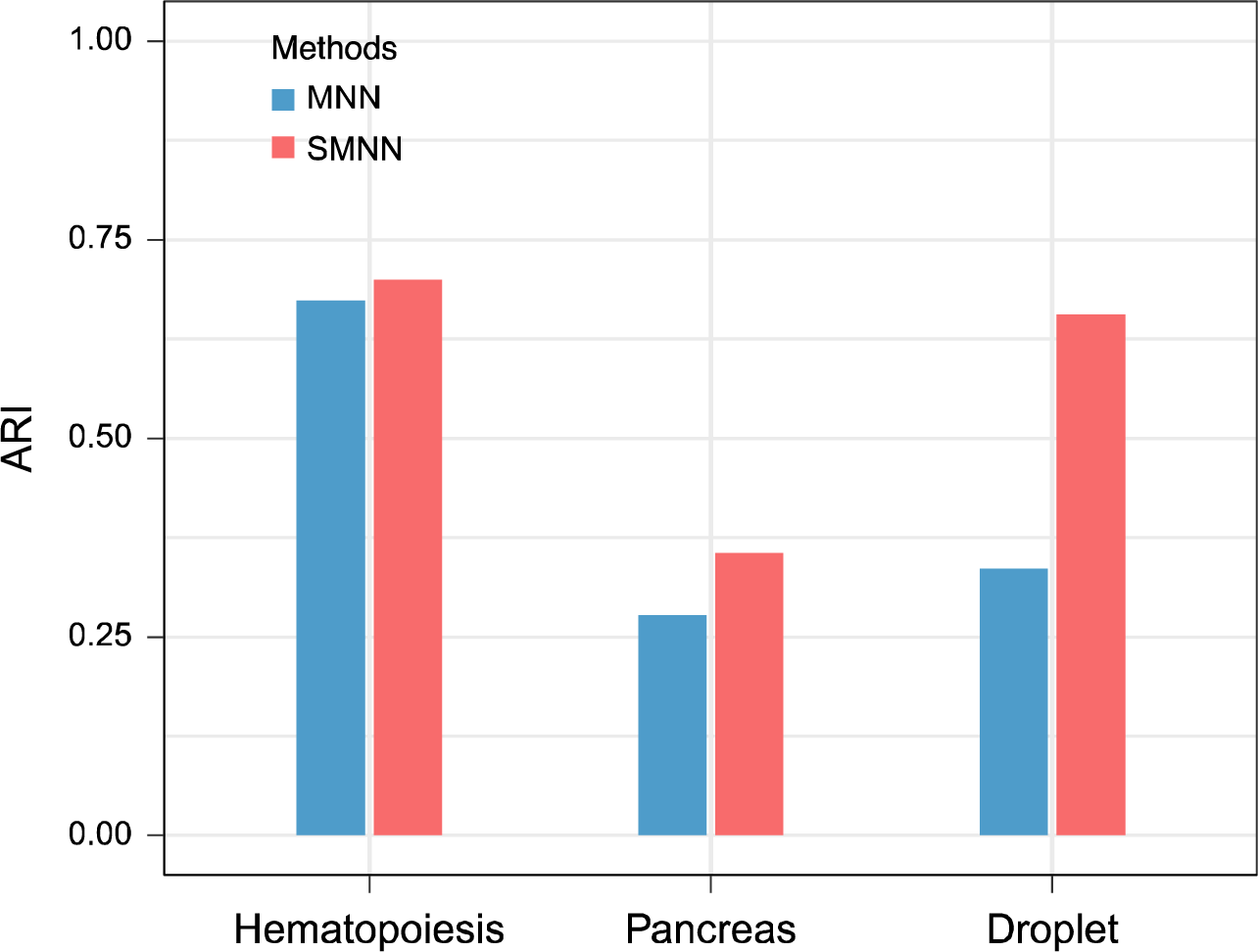
Clustering accuracy in three datasets after batch effect correction. Adjusted Rand Index (ARI) is employed to measure the similarity between clustering results before and after batch effect correction.

## Discussion

In this study, we present SMNN, a batch effect correction method for scRNA-seq data via supervised mutual nearest neighbor detection. Our work is built on the recently developed method MNN, which has showed advantages in batch effect correction than existing alternative methods. On top of MNN, our SMNN relaxes a strong assumption that underlies MNN: that the biological differentiations are orthogonal to batch effects [8]. When this fundamental assumption is violated, especially under the realistic scenario that the two batches are rather different, MNN tends to err when searching nearest neighbors for cells belonging to the same biological cell type across batches. Our SMNN, in contrast, explicitly considers cell type label information to perform supervised mutual nearest neighbor matching, thus empowered to extract only desired neighbors from the same cell type.

A notable feature of our SMNN is that it can detect and match the corresponding cell populations across batches with the help of feature markers provided by users. SMNN performs clustering within each batch before merging across batches, which can reveal basic data structure, i.e. cell composition and proportions of contributing cell types, without any adverse impact due to batch effects. Cells of each cluster are labeled by leveraging their average expression levels of certain marker(s), thus enabling us to limit the mutual nearest neighbor detection within a smaller search space (i.e., only among cells of the same or similar cell type or status). This supervised approach eliminates the correction biases incurred by pairs of cells wrongly matched across cell types. We benchmarked SMNN together with three state-of-the-art batch effect correction methods, MNN, Seurat v3 and LIGER, on simulated and three published scRNA datasets. Our results clearly show the advantages of SMNN in terms removing batch effects. For example, our results for the hematopoietic datasets show that SMNN better mixed cells of all the three cell types across the two batches (**Fig. 4a-e**), and reduced the differentiation between the two batches by up to 96.6% on top of the corrected results from the three unsupervised methods (**Fig. 4f**), demonstrating that our SMNN method can more effectively mitigate batch effect. Additionally, cell population composition also can be a critical factor in batch effect correction. Our results by analyzing batches with varying cell type compositions (detailed in **Supplementary Section 3**; **Supplementary Fig. S18**) suggest that our SMNN is robust to differential cell composition across batches.

More importantly, the wrongly matched cell pairs may wipe out the distinguishing features of cell types. This is mainly because, for a pair of cells from two different cell types, the true biological differentiations between them would be considered as technical biases and subsequently removed in the correction process. Compared to MNN, SMNN also appears to more accurately recover cell-type specific features: clustering accuracy using SMNN-corrected data increases substantially in all the three real datasets (by 7.6 to 42.3% when measured by ARI) (**Fig. 7**). Furthermore, we observe power enhancement in detecting DEGs between different cell types in the data after SMNN correction than MNN (**Fig. 5 and 6 and Supplementary Fig. S11-15**). Specifically, the precision of the DEGs identified by SMNN were improved by up to 841% and 54.0% than those by MNN when compared to the two set of working truth, respectively (**Fig. 6c and d and Supplementary Fig. S14-15**). Moreover, GO term enrichment results show that, the up-regulated DEGs identified only in SMNN-corrected GMP and MEP cells were involved in immune process and blood coagulation, respectively (**Supplementary Fig. S11f and h**), which accurately reflect the major features of these two cell types [21]. Similarly, DEGs identified between T and B cells after SMNN correction are also biologically more relevant than those identified after MNN correction (**Supplementary Fig. S17f and h**). These results suggest that SMNN can eliminate the overcorrection between different cell types and thus maintains more biological features in corrected data than MNN. Efficient removal of batch effects at reduced cost of biological information loss, manifested by SMNN in our extensive simulated and real data evaluations, empowers valid and more powerful downstream analysis.

In summary, extensive simulation and real data benchmarking suggest that our SMNN can not only better rescue biological features and thereof provide improved cluster results, but also facilitate the identification of biologically relevant DEGs. Therefore, we anticipate that our SMNN is valuable for integrated analysis of multiple scRNA-seq datasets, accelerating genetic studies involving single-cell dynamics.

## Materials and methods

### Simulation Framework

We simulated two scenarios, orthogonal and non-orthogonal, to compare the performance of MNN and SMNN. The difference between the two scenarios lies in the directions of the true underlying batch effect vectors with respect to those of the biological effects.

### Baseline simulation

Our baseline simulation framework, similar to that adopted in Haghverdi *et al*. [8], contains two steps:

First, data are initially generated in low (specifically three) dimensional biological space. Data in each batch are independently generated from a Gaussian mixture model to represent a low dimensional biological space, with each component in the mixture corresponding to one cell type. Equations (1) and (2) below show formulae to generate two batches of such initial data, represented by matrices X_*k*_ and Y_*l*_, in low dimensional biological space.

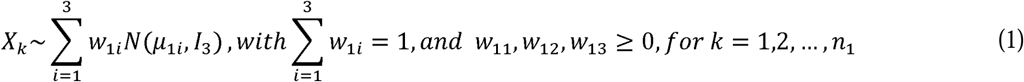

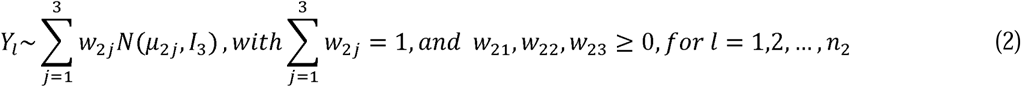

where *μ*_1*i*_ is the three-dimensional vector specifying cell-type specific means for the *i*-th cell type in the first batch, reflecting the biological effect; similarly for *μ*_2*j*_; *n*_1_ and *n*_2_ is the total number of cells in the first and second batch, respectively; *w*_1*i*_ and *w*_2*j*_ are the different mixing coefficients for the three cell types in the two batches; and *I*_3_ is the three dimensional identity matrix with diagonal entries as ones and the rest entries as zeros. In our simulations, we set *n*_1_ = 1000, *n*_2_ = 1100 and

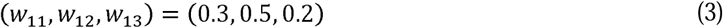

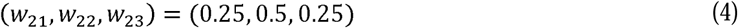

Secondly, we project the low dimensional data with batch effect to the high dimensional gene expression space. We map both datasets to *G* = 50 dimensions by linear transformation using the same random Gaussian matrix ***P***, to simulate high-dimensional gene expression profiles.

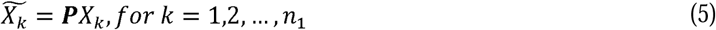

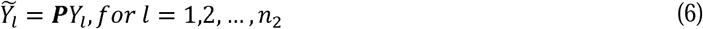

Here ***P*** is a *G*×3 Gaussian random matrix with each entry simulated from the standard normal distribution.

### Introduction of batch effects

In Haghverdi *et al*. [8], batch effects are directly introduced in the high dimensional gene expression space. Specifically, a Gaussian random vector *b* = (*b*_1_,*b*_2_,…,*b*_*G*_)^*T*^ is simulated and added to the second dataset via the following:

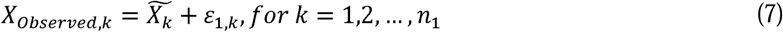

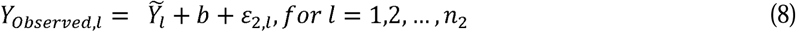

where 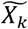 and 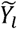 are projected high-dimensional gene expression profiles; *ε*_1,*k*_ and *ε*_2,*l*_ are independent random noises added to the expression of each “gene” for each cell in the two batches.

In our simulations, we adopt a slightly different approach: we introduce batch effects in the low dimensional biological space. Specifically, we simulate a bias vector *c* = (*c*_1_,*c*_2_,*c*_3_)^*T*^ in the biological space:

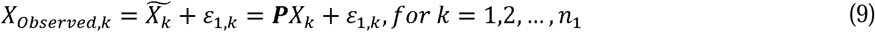

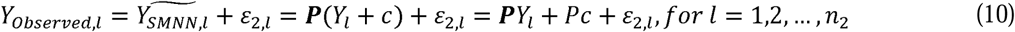

Our simulation framework can be viewed as a reparametrized version of the model in Haghverdi *et al*. [8]. For each batch effect ***b*** of the model in Haghverdi *et al*. [8], there exist multiple pairs of projection matrix ***P*** and vector *c* such that *b* = ***P****c*, and for any vector *c* in our model, there is a corresponding vector *b* = ***P****c* given a fixed projection matrix ***P***. In particular, 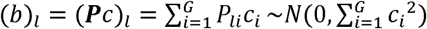, if 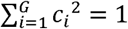. In other words, for any simulated setting in Haghverdi *et al*. [8], we can find at least one equivalent setting in our model; and vice versa. Although our simulation framework is largely similar to that in Haghverdi *et al*. [8], the two differ in the following two aspects:

First, the lowdimensional biological space is three-dimensional in ours and two-dimensional in Haghverdi *et al*. [8].

Second, we introduce batch effects *c* in low dimensional biological space and then projected to high dimensional space (equation (10)), while Haghverdi *et al*. [8] directly introduce batch effects *b* in the high dimensional gene expression space (equation (8)). We made such changes so that we can simulate both the orthogonal and non-orthogonal scenarios in a more straightforward manner the extent of orthogonality can be controlled (equation (11)). The orthogonality is defined in the sense that biological differences (that is, mean difference between any two clusters/cell-types), are orthogonal to those from batch effects.

Our framework allows flexible modeling of the biological effects and batch effects in the same low dimensional biological space and allow us to control the extent of orthogonality. Specifically, the batch effect *c* is added to mean vectors of three cell types in batch 1 to get the mean vectors of three cell types for batch 2.

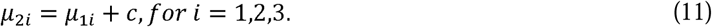

Note that (*μ*_1*j*_ − *μ*_1*i*_)*c* = 0, *for i* ≠ *j* ∈ {1,2,3} represents the orthogonal scenario that variation from batch effect is orthogonal to mean difference between any two clusters/cell-types, and (*μ*_1*j*_ − *μ*_1*i*_)*c* ≠ 0, *for i* ≠ *j* ∈ {1,2,3} in the non-orthogonal case.

Leveraging the simulation framework described before, we simulate two scenarios via the following:

1. In the orthogonal case, we set *c* = (0, 0, 2)^*T*^
  a. *μ*_11_ = (5, 0, 0)^*T*^, *μ*_12_ = (0, 0, 0)^*T*^, *μ*_13_ = (0, 5, 0)^*T*^
  b. *μ*_21_ = (5, 0, 2)^*T*^, *μ*_22_ = (0, 0, 2)^*T*^, *μ*_23_ = (0, 5, 2)^*T*^
2. In the non-orthogonal case, we set *c* = (0,5,2)^*T*^
  a. *μ*_11_ = (5, 0, 0)^*T*^, *μ*_12_ = (0, 0, 0)^*T*^, *μ*_13_ = (0, 5, 0)^*T*^
  b. *μ*_21_ = (5, 5, 2)^*T*^, *μ*_22_ = (0, 5, 2)^*T*^, *μ*_23_ = (0, 10, 2)^*T*^

### Performance evaluation

MNN and SMNN share the goal to correct batch effects. Mathematically, using the notations introduced in baseline simulation, the goal translates into de-biasing vector *c* (which would be effectively reduced to *b* in the orthogonal case). Without loss of generality and following MNN, we treat the first batch as the reference and correct the second batch {*Y*_*Observed,l*_: *l* = 1, …, *n*_2_} to the first batch {*X*_*Observed,k*_: *k* = 1, …, *n*_1_}. Denote the corrected values from MNN and SMNN as 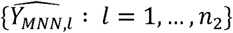 and 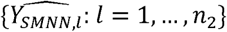, respectively.

To measure the performance of the two correction methods, we utilize the Frobenius norm [20] to define the loss function:

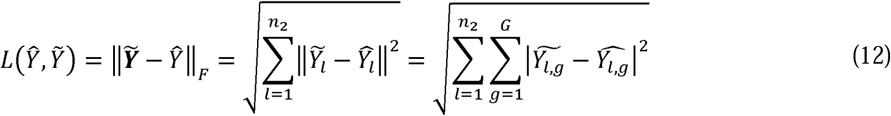

where 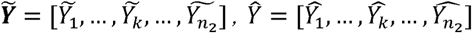. Note that 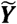, is the simulated true profiles introduced in equations (5) and (6) before batch effects, and noises are introduced in equations (7) and (8). Since MNN conducts cosine normalization to the input and the output, we use cosine-normalized 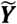 when calculating the above loss function.

### Real data benchmarking

To assess the performance of SMNN in real data, we compared SMNN to alternative batch effect correction methods: MNN [8], Seurat v3 [17], and LIGER [22] to two hematopoietic scRNA-seq datasets, generated using different sequencing platforms, MARS-seq and SMART-seq2 (**Supplementary Table S1**) [10, 23]. The first batch produced by MARS-seq consists of 1920 cells of six major cell types, and the second batch generated by SMART-seq2 contains 2730 of three cell types, where three cell types, CMP, GMP and MEP cells, are shared between these two batches (here the two datasets). Batch effect correction was carried out using all four methods, following their default instructions. Cell type labels were fed to SMNN directly according to the annotation from the original papers. To better compare the performance between MNN and SMNN, only the three cell types shared between the two batches were extracted for our downstream analyses. The corrected results of all the three cell types together, as well as for each of them separately, were visualized by UMAP using *umap-learn* method [24]. In order to qualify the mixture of single cells using both batch correction methods, we calculated: 1) F statistics under two-way multivariate analysis of variance (MANOVA) for merged datasets of the two batches. F statistics quantifies differences between batches, where smaller values indicating better mixing of cells across batches; and 2) the distance for the cells within each cell type in batch 2 to the centroid of the corresponding cell group in batch 1.

To measure the separation of cell types after correction, we additionally attempted to detect DEGs between different cell types in both SMNN and MNN corrected datasets. The corrected expression matrices of the two batches were merged and DEGs were detected by Seurat v3 using Wilcoxon rank sum test [17]. Genes with an adjusted *p*-value < 0.01 were considered as differentially expressed. GO enrichment analysis was performed for the DEGs exclusively identified by SMNN using *clusterProfiler* [25]. Because there is no ground truth for DEGs, we further identified DEGs between different cell types within corrected batch 2 and then compared them to those identified in uncorrected batch 1 and uncorrected batch 2, which supposedly are not affected by the choice of batch effect correction method. TPR was computed for each comparison.

Additionally, we also performed batch effect correction on another two tissues/cell lines, pancreas [26, 27] and PBMCs [28], again using both SMNN and MNN. DEGs were detected between T cells and B cells in the merged PBMC and T cell datasets after SMNN and MNN correction, respectively. Furthermore, single cell clustering was applied to batch-effects corrected gene expression matrices in all the three real datasets following the pipeline described in Haghverdi *et al*. [8]. Cell type labels before correction were considered as ground truth and ARI [29] was employed to measure the clustering similarity before and after correction:

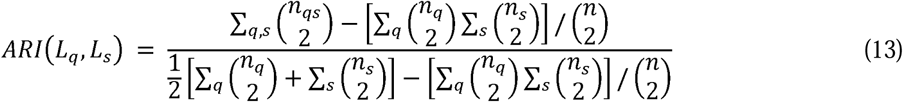

where *n*_*q*_ and *n*_*s*_ are the single cell numbers in cluster *q* and *s*, respectively; *n*_*qs*_ is the number of single cells shared between clusters *q* and *s*; and *n* is the total number of single cells. ARI ranges from 0 to 1, where a higher value represents a higher level of similarity between the two sets of cluster labels.

## Data and software availability

SMNN is compiled as an R package, and freely available at https://yunliweb.its.unc.edu/SMNN/ and https://github.com/yycunc/SMNN. The data we adopted for benchmarking at from following: 1) two Mouse hematopoietic scRNA-seq datasets from [10] (GEO accession number GSE81682) and [23] (GEO accession number GSE72857); 2) two human pancreas scRNA-seq datasets from [26] (GSE81076) and [27] (GSE85241); and two 10X Genomics datasets of PBMCs and T cells from [28] (https://support.10xgenomics.com/single-cell-gene-expression/datasets/).

## Supplementary Data

Supplementary materials are available online.

## Author Contributions

Y.L. initiated and designed the study. Y.Y., G.L. and H.Q. implemented the model and performed simulation studies and benchmarking evaluation. Y.Y, G.L. and Y.L. wrote the manuscript and all authors edited and revised the manuscript.

