## Supplementary materials for "SMNN: Batch Effect Correction for Single-cell RNA-seq data via Supervised Mutual Nearest Neighbor Detection"

**Section 1. Outline of our SMNN framework**

Perform unsupervised clustering using Seurat for each batch, from ${batch}_{1}$to ${batch}_{n}$. // Users can skip this step by feeding SMNN cell type labels that are consistent or comparable across batches.

**For** ${batch}_{1}$ to ${batch}_{n}$**DO**

Compute the average expression value of each marker gene, for each cluster identified in this batch

Compute the proportion of cells of each cluster expressing each of the marker genes

Assign labels to the cluster(s) that meet both two criteria above

**End**

**For** $i=2$ to $n$**DO** // ${batch}_{1}$ serves as the reference batch and would be returned in the output as is without correction

Detect mutual nearest neighbors within each matched cell type between ${batch}_{1}$ and ${batch}_{i}$

Compute batch effect correction vector for each identified pair of cells

Compute the cell-specific correction vectors by exploiting a Gaussian kernel

Scale the correction factors based on

1) the cell’s locations on the correction vector space and

2) the quantiles between batches

to ensure that their quantiles after correction are equal. // To eliminate “kissing effects”

**Return** 1) Batch-effects corrected expression matrix for ${batch}_{i}$

2) Information regarding nearest neighbors between ${batch}_{1}$ and ${batch}_{i}$

**End**

**Section 2. Correction for the cells in batch-specific clusters**

The correction vector obtained from shared cell-types will be applied to correct all the cells, including the cells in batch-specific clusters. Specifically, for each cell x in the target batch (which will be corrected), whether it falls into a batch-shared cluster or a batch-specific cluster, SMNN calculates a cell-specific correction vector using the same formula below. The cell-specific correction vector $\vec{u}$ for any target cell x is a weighted sum of vector differences $\vec{v}_{ml}$ across all identified MNN pairs (without loss of generosity, we assume m is from the target batch and l from the reference batch. Note that in SMNN, the two cells m and l in each MNN pair belong to the same cluster/cell-type across the two batches).

$$\vec{u}(x)=\frac{\sum_{MNN pairs} \vec{v}_{ml}W\left( x,m \right)}{\sum_{MNN pairs} W\left( x,m \right)}$$

The Gaussian kernel weights$W\left( x,m \right)$ gives higher weights to vector differences ($\vec{v}_{ml}$’s) involving a cell m in closer proximity with the target cell x. The smaller the distance, the larger of the weight. Note that a cell m may appear multiple times and contributes to multiple terms in the weighted sum if it has multiple MNNs identified by our SMNN. Also note that if cell x belongs to a batch-specific cluster, itself would not contribute to any of the terms because it would not have any MNNs.

**Section 3. Additional performance evaluation**

In this study, we further evaluated several other computing and performance aspects of SMNN. First, we evaluated SMNN’s performance in eliminating the “kissing effect” introduced during batch effect correction. SMNN was applied to the two hematopoietic datasets with or without attempts to avoid “kissing effects” (by standardizing according to quantiles across batches or not). We then visualized both sets of corrected data, as well as uncorrected data, in a three-dimensional UMAP space. The results show that without standardizing, we observe quite apparent “kissing effect”: only the surfaces of the cell clouds from the two batches were brought into contact (more pronounced for MEP cells). In contrast, cell clouds across batches were merged beyond surface level after standardizing (**Supplementary Fig. S1**). Therefore, SMNN by default applies the standardization to eliminate the “kissing effect” during batch effect correction.

Secondly, we evaluated SMNN’s robustness to the availability of only partial cell type information. Specifically, we performed SMNN correction for two hematopoietic datasets by feeding cluster labels for either one or two of the three cell types. The results showed that, under such partial label information, SMNN still better mixed cells of the same cell type across batches than MNN method (**Supplementary Fig. S10a-c and e-g**), consistent with the best/lowest F values from the SMNN-corrected results F value, compared with uncorrected and MNN-corrected data. Specifically, SMNN reduced the differentiation between the two batches by 89.7% to 94.0% (73.9% to 96.7%) on top of the MNN corrected results when using label information for only one (two) of the three cell types, respectively (**Supplementary Fig. S10d and h**). These results indicate that SMNN is robust and superior even with partial cell type annotation information.

Furthermore, to assess whether the differentially expressed genes (DEGs) identified in SMNN and MNN corrected data were sensitive to the significance threshold selected, we show ROC curves to illustrate the ability of identifying DEGs among CMP, GMP and MEP cells after SMNN and MNN correction, at various adjusted p-value threshold, treating those identified in uncorrected batch 1 as working truth. The results showed that, in almost all cases, SMNN outperformed MNN (**Supplementary Fig. S16**). The only exception is for DEGs up-regulated in GMP when compared to MEP, where the ROC curves of SMNN and MNN are comparable (**Supplementary Fig. S16e**). These results suggest that our DEG analysis is not sensitive to the *p*-value cutoff we use for DEG detection.

Lastly, when the cell population compositions are unbalanced between two batches, we extracted the three major cell types out from two hematopoietic datasets, in order to construct two new batches with a more apparent difference in cell group composition. In the new batch 1, the proportion of CMP, GMP and MEP cells are 40.4%, 15.1% and 44.5%, while those are 17.6%, 42.3% and 40.1% in new batch 2. Then we performed both MNN and SMNN correction across the new batches. The results showed that SMNN outperformed MNN that SMNN reduced the differentiation between the two batches (measured by the F value) by 11.3% on top of the MNN corrected results (**Supplementary Fig. S18**), indicating that SMNN is robust to the cell population composition.

**Table S1.** Major characteristics of the three benchmarking datasets.

| **Dataset** | **Batch ID** | **Tissue origin** | **# of cells** | **Technical platform** | **Ref** |
| --- | --- | --- | --- | --- | --- |
| Hematopoiesis | batch 1 | Mouse hematopoiesis | 1,920 | SMART-seq2 | Nestorowa *et al*. 2016 |
|  | batch 2 | Mouse hematopoiesis | 2,730 | MARS-seq | Paul *et al*. 2015 |
| Pancreas | batch 1 | Human cadaveric pancreata | 1,007 | CEL-seq | Grün *et al.* 2016 |
|  | batch 2 | Human cadaveric pancreata | 1,595 | CEL-seq2 | Muraro *et al.* 2016 |
| 10X Genomics | batch 1 | PBMC | 68,580 | 10X Genomics GemCode | Zheng et al. 2017 |
|  | batch 2 | T cells | 4,459 | 10X Genomics GemCode | Zheng et al. 2017 |

**Supplementary Figures:**

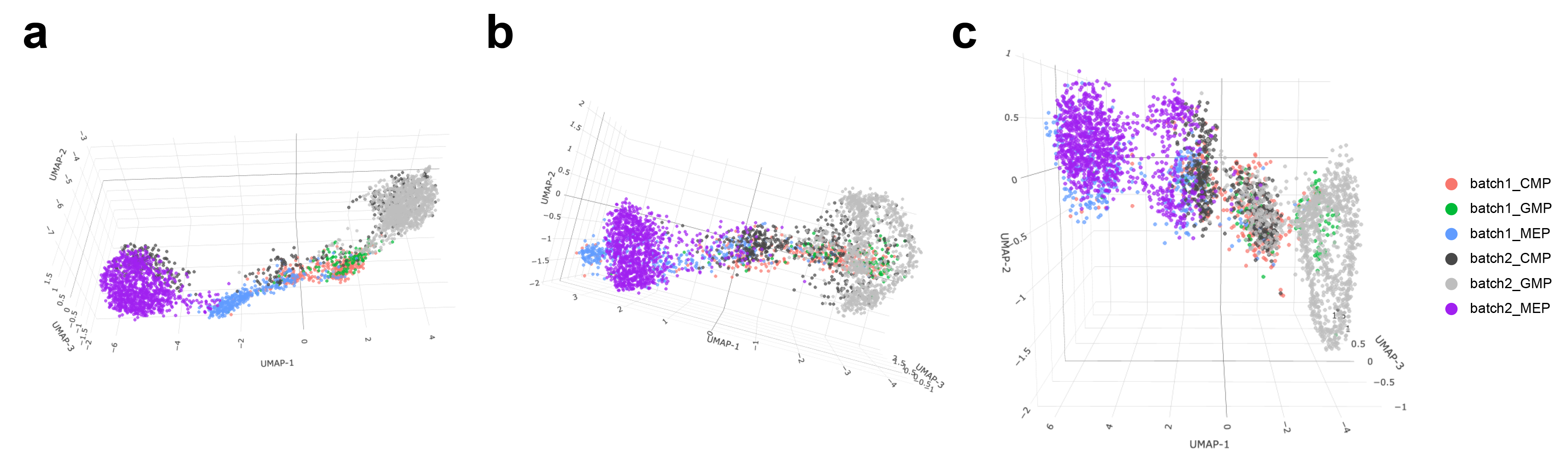

**Fig. S1. Performance of SMNN in eliminating “kissing effects”. (a)** 3D UMAP plot for the two hematopoietic datasets before batch effect correction. Different cell types in different batches are shown in different colors. (**b**) 3D UMAP plot without adjusting “kissing effects”. (**c**) 3D UMAP plot after adjusting “kissing effects”.

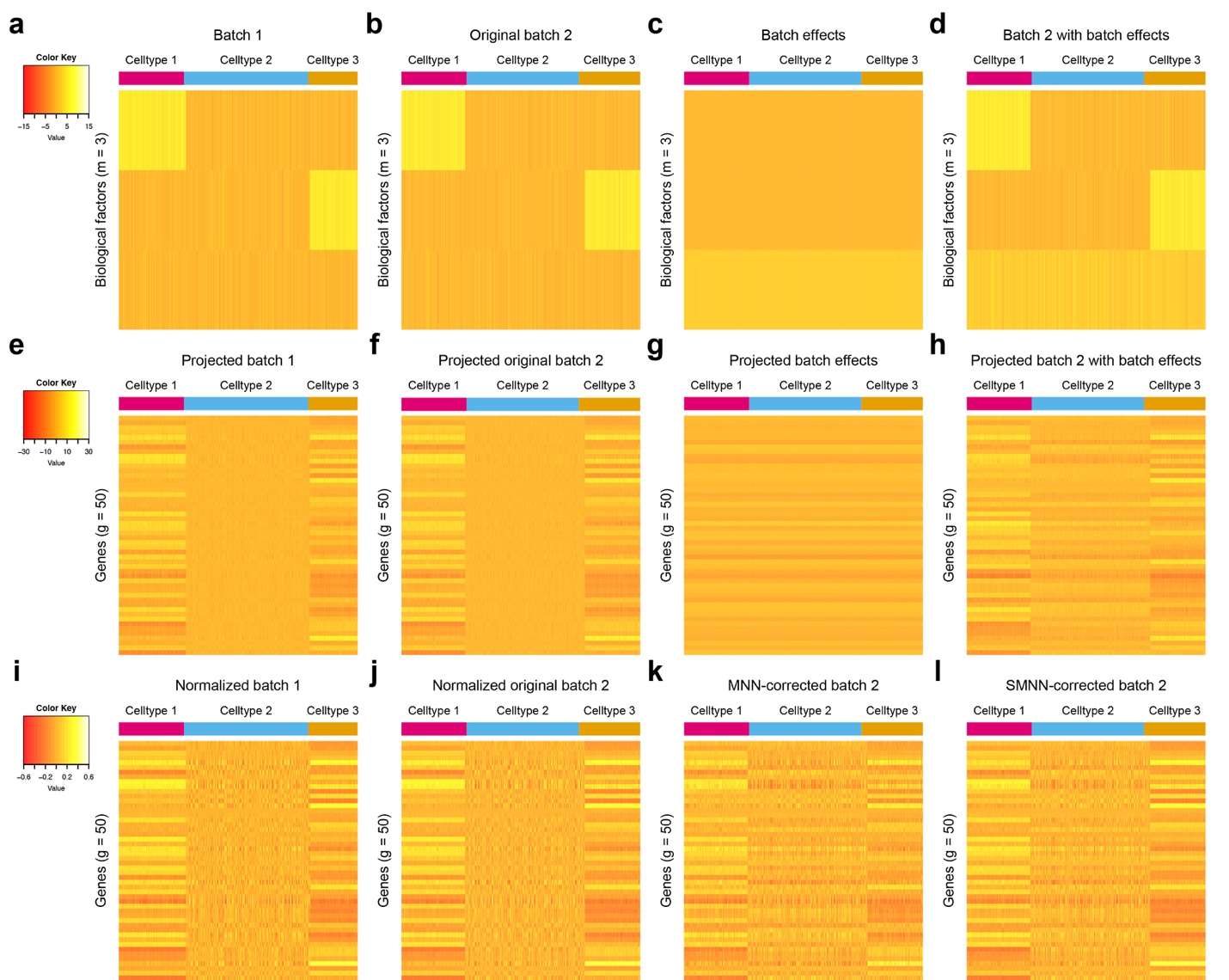

**Fig. S2.** **Heatmap matrix for simulated data under the orthogonal scenario.** (**a**), (**b**), (**c**) and (**d**) show the 3-dimensional biological space with rows of each heatmap representing biological factors and columns corresponding to single cells. (**e**), (**f**), (**g**) and (**h**) show the high dimensional gene expression profiles with rows corresponding to genes and columns again corresponding to single cells. (**a**), (**e**) and (**i**) correspond to the batch 1, and (**b**), (**f**) and (**j**) correspond to batch 2. (**c**) and (**g**) provide a visualization for the direction of batch effects in low-dimension biological space and high-dimension gene expression spaces, respectively. (**d**) and (**h**), sum of (**b**) and (**c**) and sum of (**f**) and (**g**) respectively, are “observed” data for cells in batch 2 in low and high dimensional space respectively. (**i**) and (**j**) are the cosine-normalized data for batch 1 and original batch 2. Note “original” is in the sense that no batch effects have been introduced to the data yet. (**k**) and (**l**) are the MNN and SMNN corrected results.

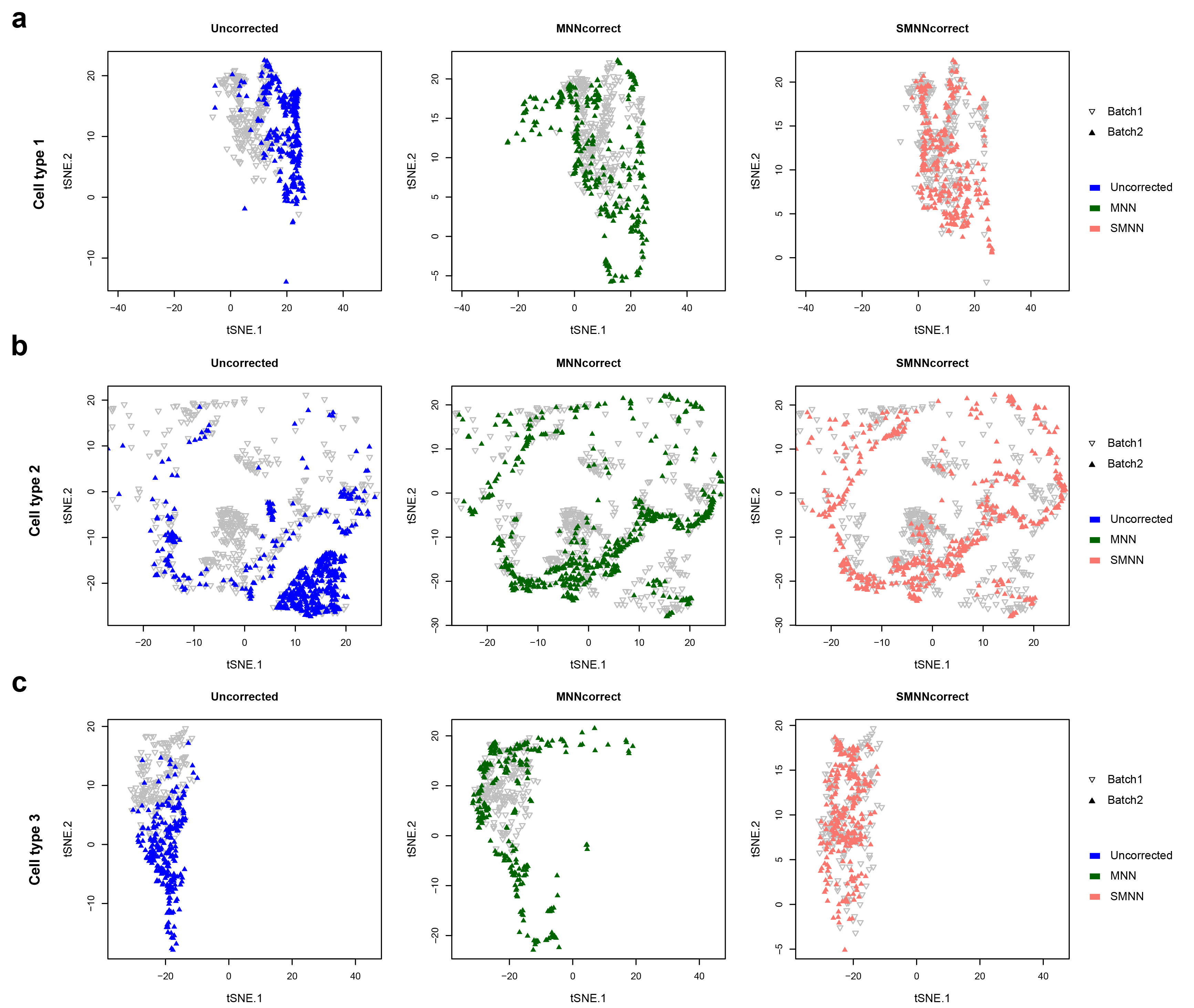

**Fig. S3.** **t-SNE plots by cell type under the orthogonal scenario.** The first column shows the uncorrected data after cosine normalization; the second column shows the MNN corrected results and the last column shows the SMNN corrected results. (**a**), (**b**) and (**c**) correspond to three different cell types.

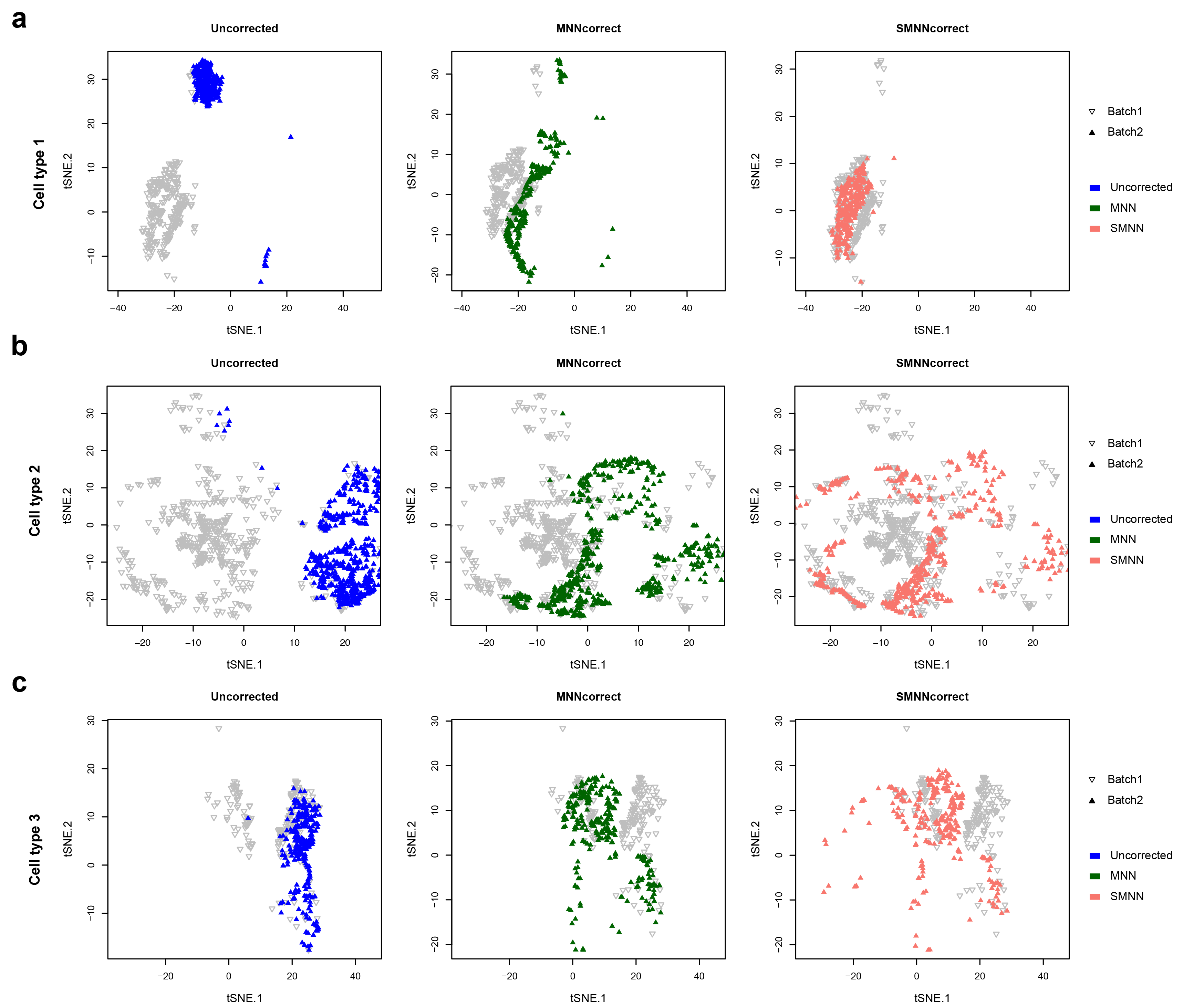

**Fig. S4. t-SNE plots by cell type under non-orthogonal scenario.** The first column shows the uncorrected data after cosine normalization; the second column shows the MNN corrected results and the last column shows the SMNN corrected results. (**a**), (**b**) and (**c**) corresponds to three different cell types.

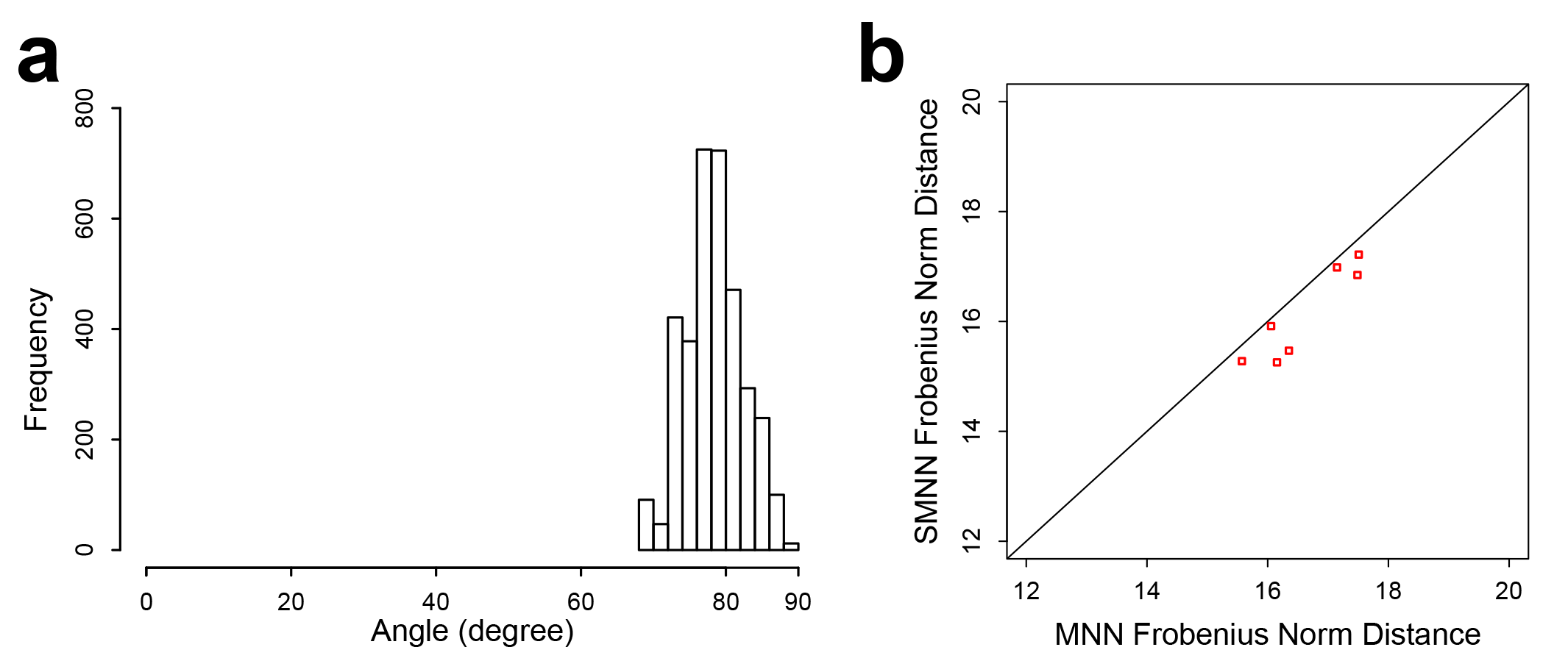

**Fig. S5. Performance of SMNN and MNN in the simulation data using the model in Haghverdi *et al.* (2018)**. (**a**) Histogram of the angles (surrogate for orthogonality) for the simulation data using the model in Haghverdi *et al.* (2018). (**b**) Frobenius norm distance between two batches after SMNN and MNN correction in simulation data under orthogonal (left) and non-orthogonal scenarios (right).

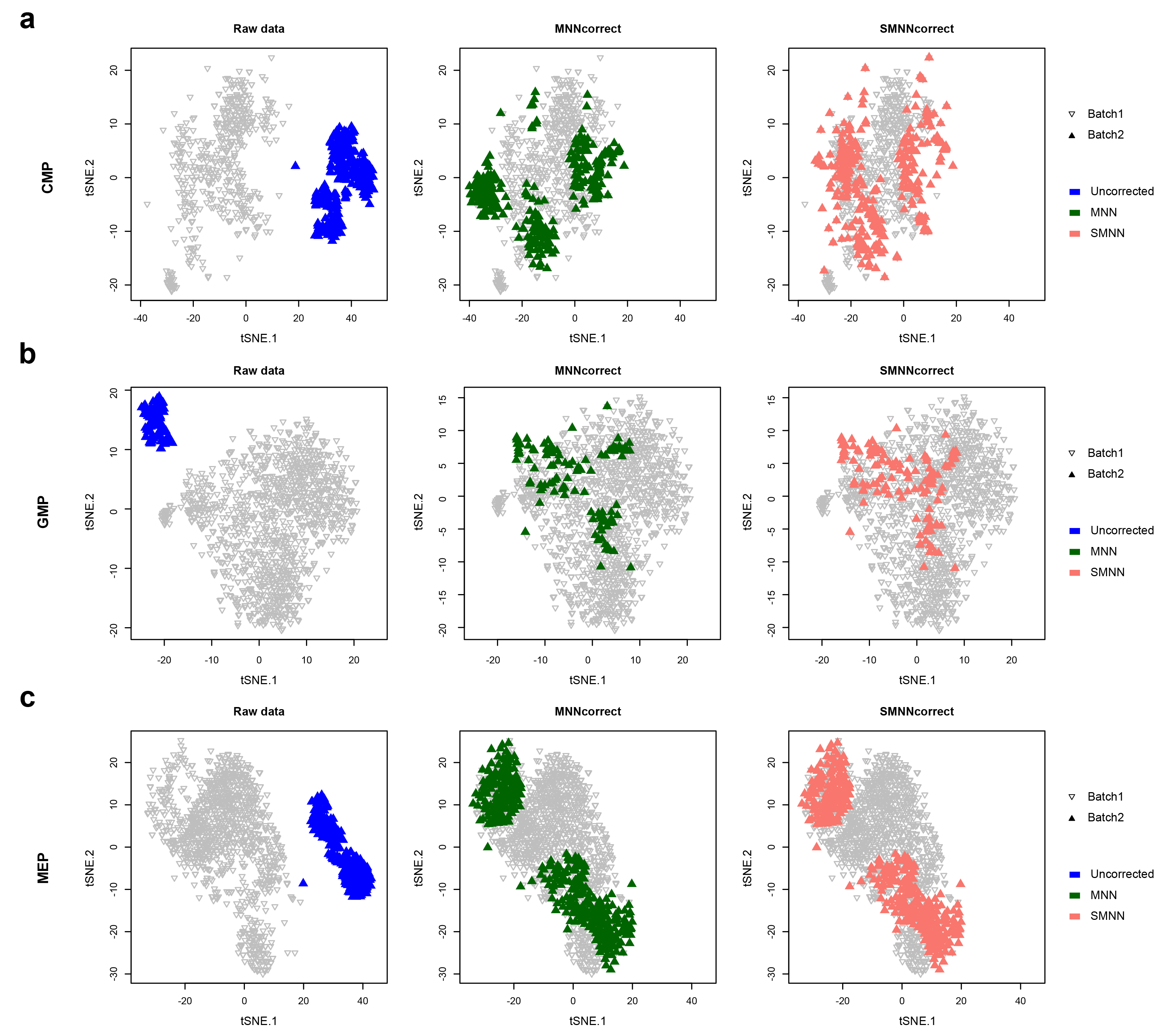

**Fig. S6. t-SNE plots by cell type in two hematopoietic datasets.** (**a**), (**b**) and (**c**) corresponds to three different cell types, CMP, GMP and MEP. The first column shows the raw data; the second column shows the MNN corrected results and the last column shows the SMNN corrected results.

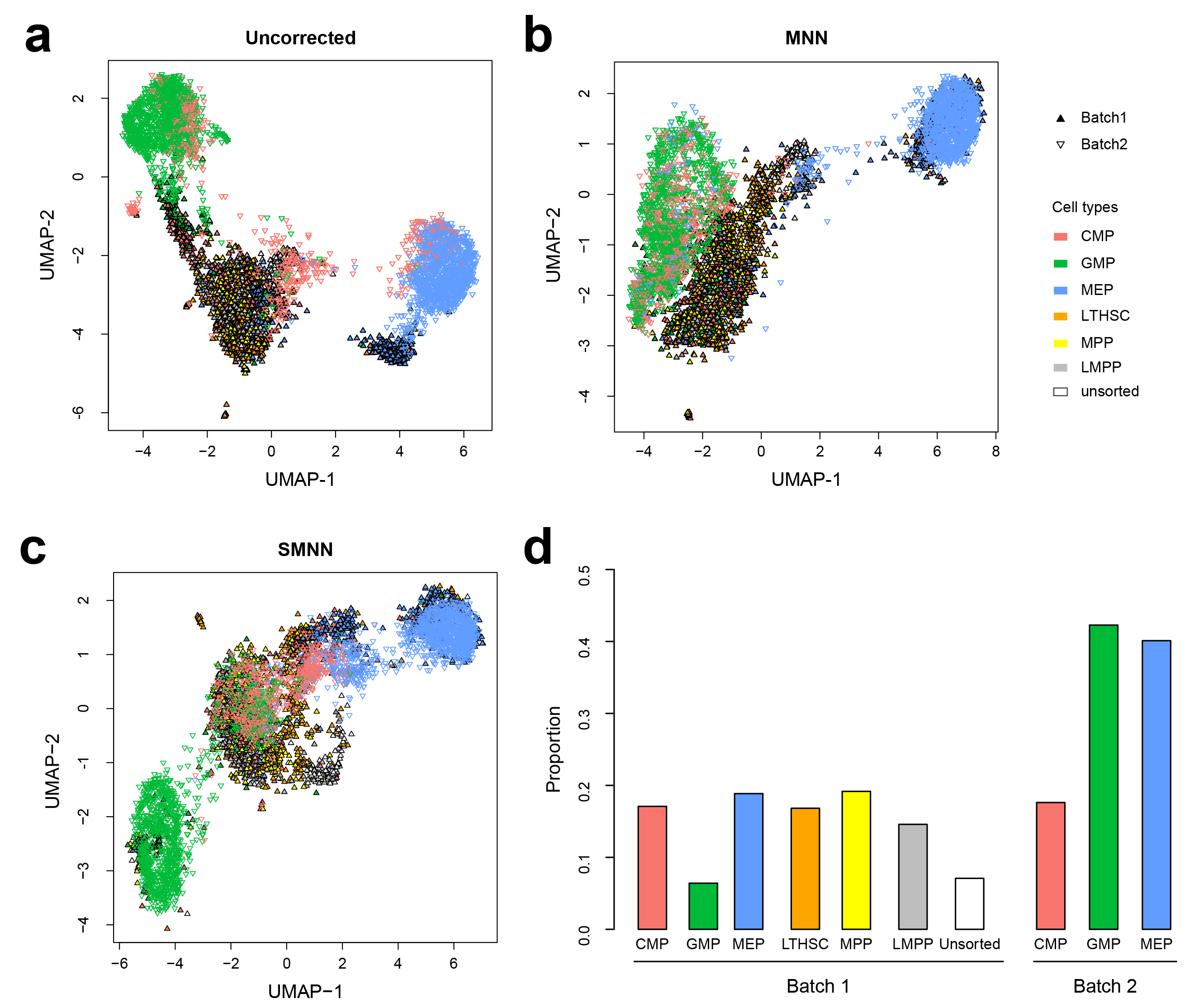

**Fig. S7.** **UMAP plots for all the cell types in two hematopoietic datasets.** (**a**), (**b**) and (**c**) corresponds to uncorrected, MNN-corrected and SMNN corrected results, respectively. (**d**) Cell type proportions in the two batches.

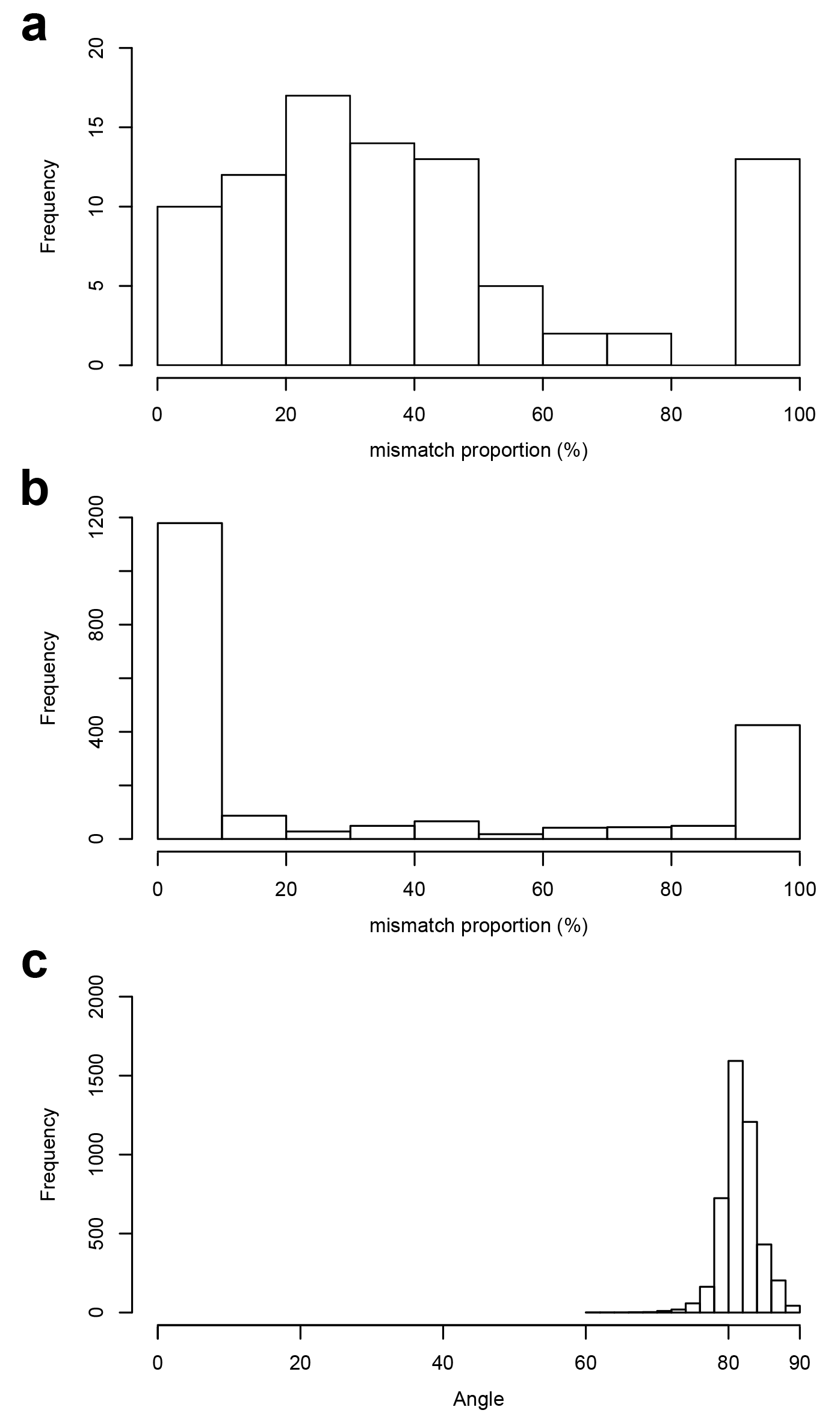

**Fig. S8. Quantification of cell-type mismatching in MNN identified nearest neighbors**. (**a**) Histogram of the proportion of nearest neighbors from a mismatching cell type (for each cell), for two hematopoietic datasets. (**b**) Histogram of the proportion of nearest neighbors from a mismatching cell type (for each cell), for the 10X Genomics datasets. (**c**) Histogram of the angles (surrogate for orthogonality) for the 10X Genomics datasets.

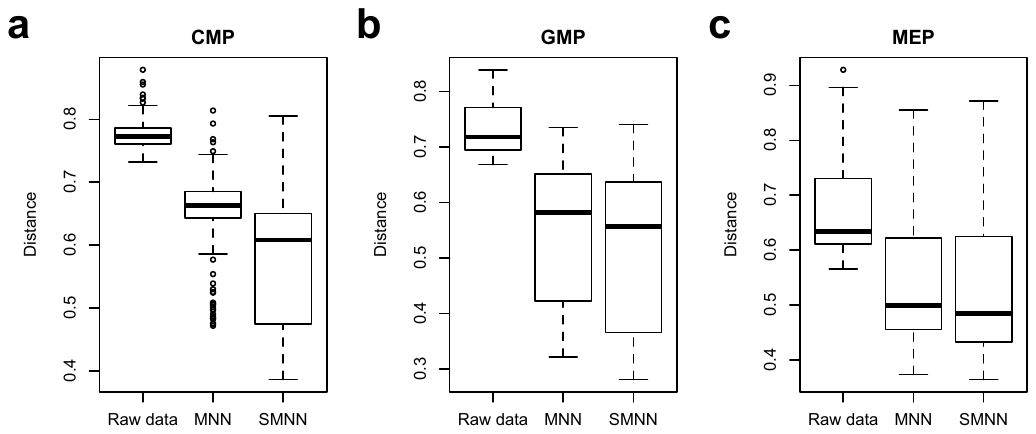
**Fig. S9.** Boxplot of distances by cell type. For each of the three different cell types, CMP (**a**), GMP (**b**) and MEP (**c**), we calculate the distance for every cell in batch 2 to its corresponding centroid in batch1.

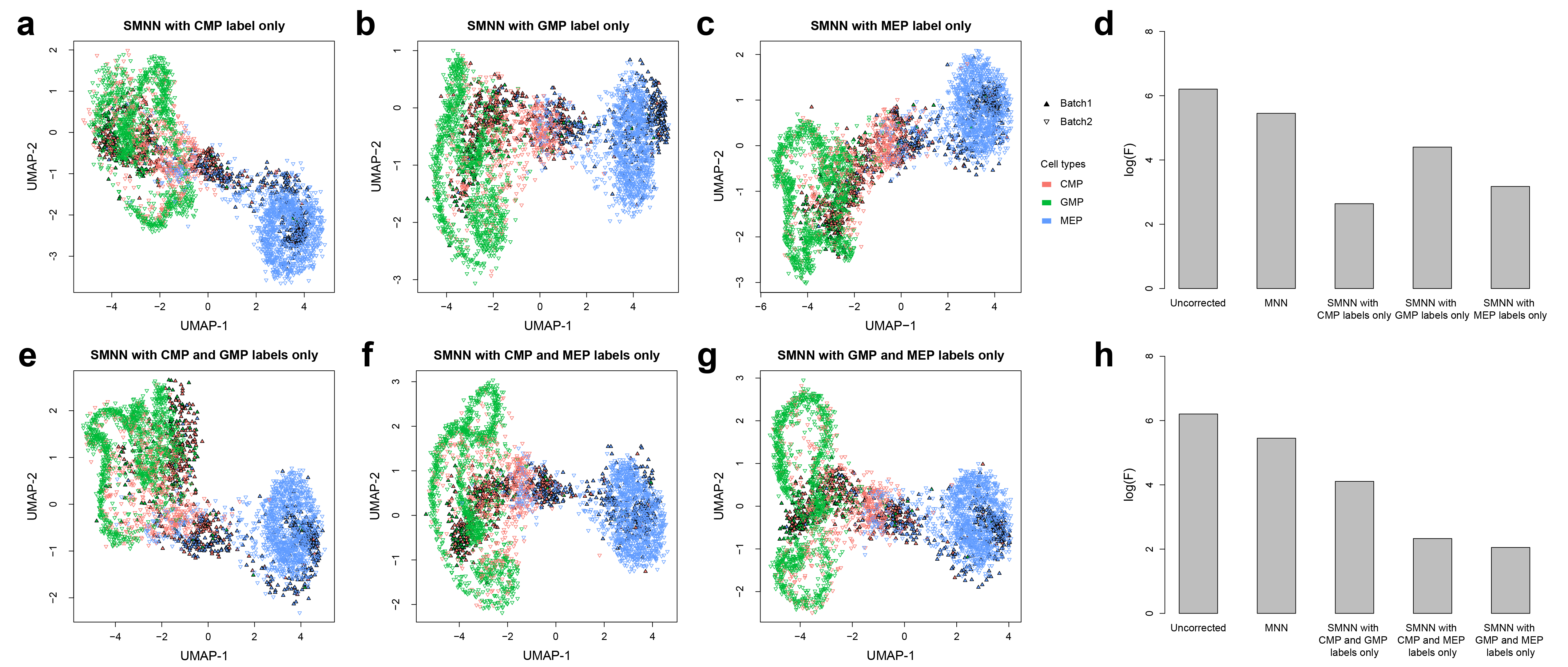

**Fig. S10. Performance of SMNN in two hematopoietic datasets with partial cell type labels.** (**a**-**c**) UMAP plots for two hematopoietic datasets after SMNN correction with the cell type label only from CMP, GMP and MEP cells, respectively. Solid and inverted triangle represent the first and second batch, respectively; and different cell types are shown in different colors. (**d**) Logarithms of F-statistics for merged data of the two batches before and after correction with MNN and SMNN. SMNN correction was performed with the cell type label only from CMP, GMP and MEP cells, respectively. (**e-g**) UMAP plots for the two hematopoietic datasets after SMNN correction with cell type label only from any two of the three cell types, CMP, GMP and MEP cells. (**h**) Logarithms of F-statistics for merged data of the two batches before and after correction with MNN and SMNN. SMNN correction was performed with the cell type label only from any of the three cell types, CMP, GMP and MEP cells.

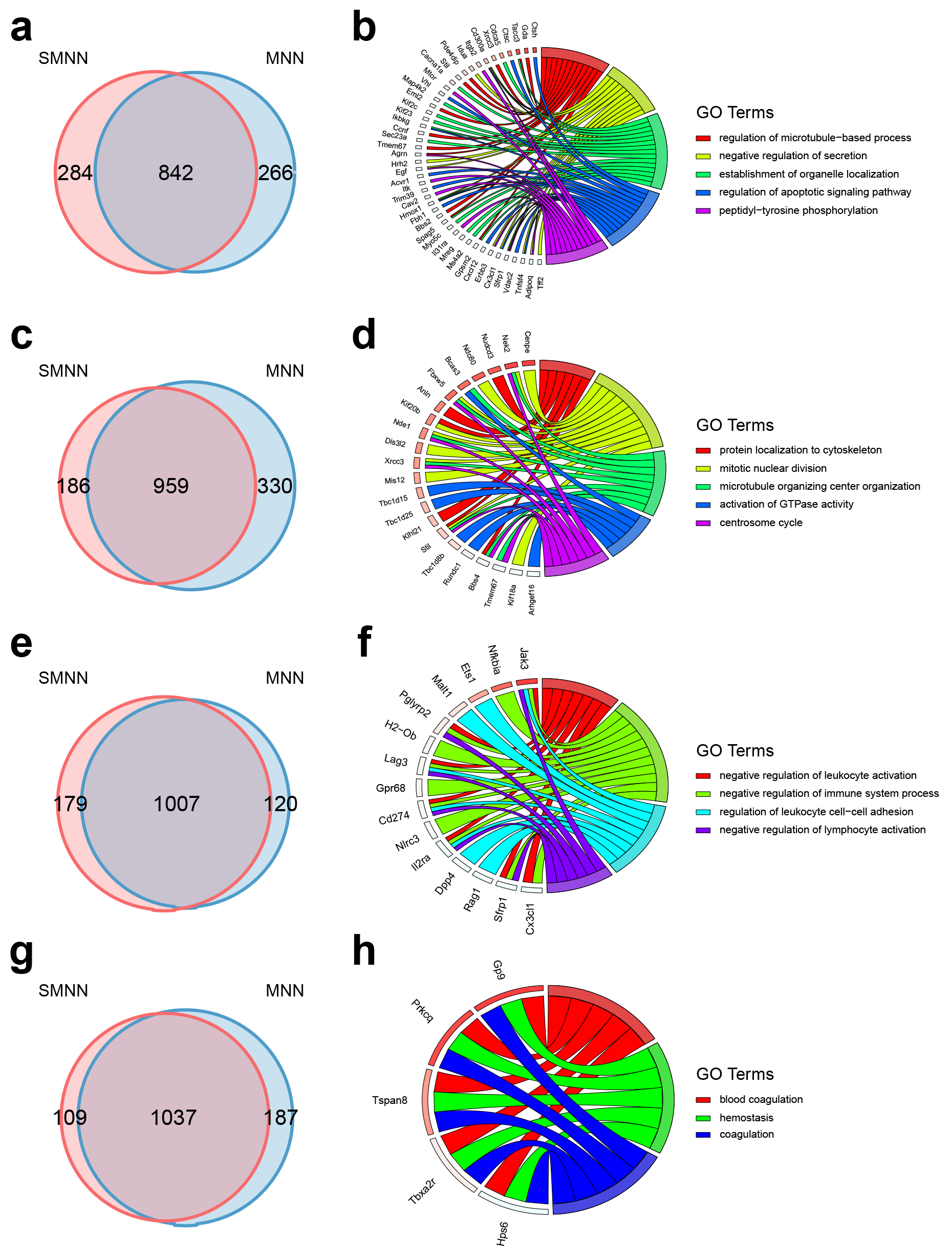

**Fig. S11. Comparison of differentially expressed genes (DEGs), identified in the merged dataset by pooling batch 1 data with batch 2 data after SMNN and MNN correction**. (**a**) Overlap of DEGs up-regulated in GMP over CMP after SMNN and MNN correction. (**b**) Feature enriched GO terms and the corresponding DEGs up-regulated in GMP over CMP. (**c**) Overlap of DEGs up-regulated in MEP over CMP after SMNN and MNN correction. (**d**) Feature enriched GO terms and the corresponding DEGs up-regulated in MEP over CMP. (**e**) Overlap of DEGs up-regulated in GMP over MEP after SMNN and MNN correction. (**f**) Feature enriched GO terms and the corresponding DEGs up-regulated in GMP over MEP. (**g**) Overlap of DEGs up-regulated in MEP over GMP after SMNN and MNN correction. (**h**) Feature enriched GO terms and the corresponding DEGs up-regulated in MEP over GMP.

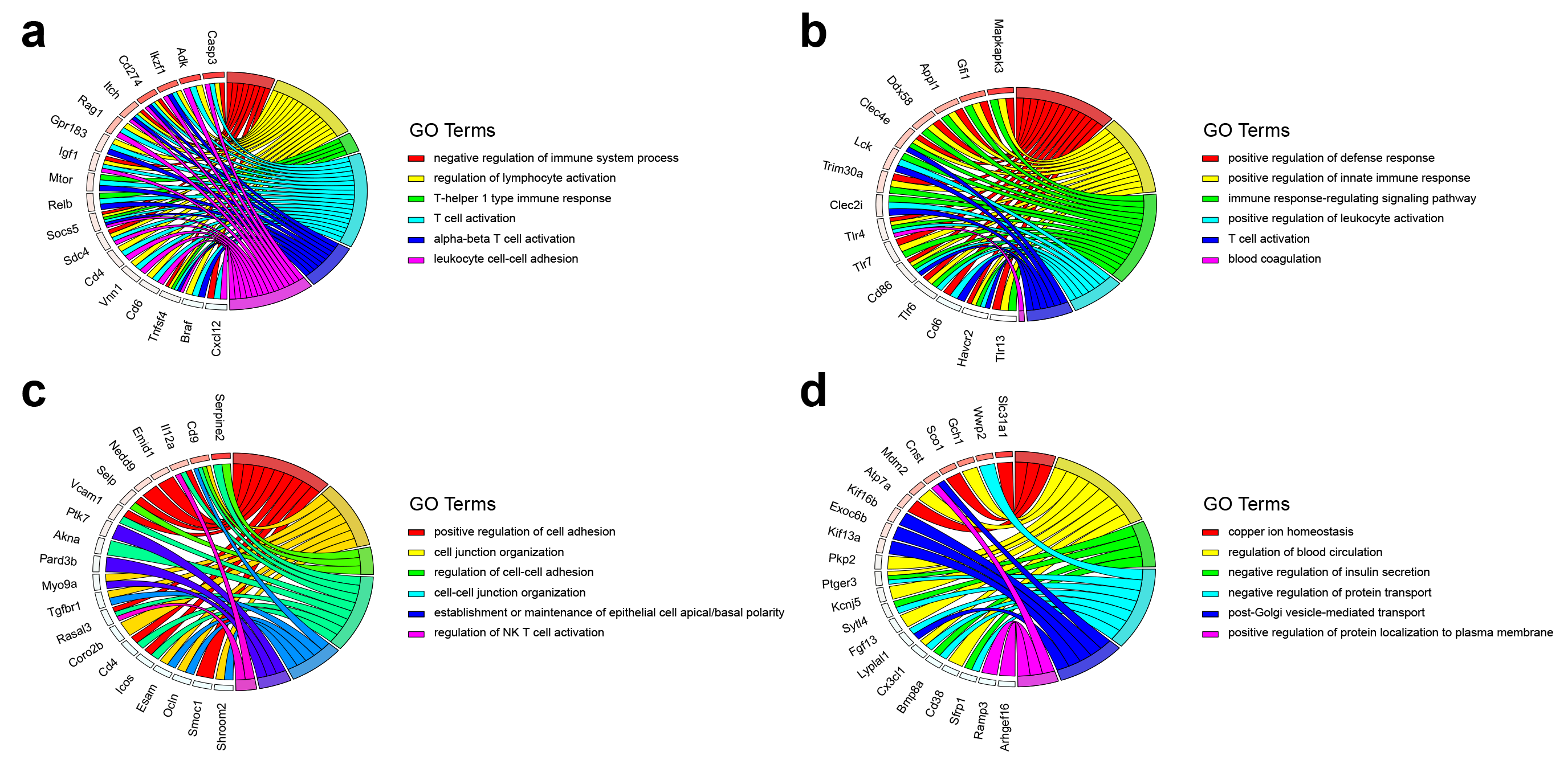

**Fig. S12. Feature enriched GO terms and the corresponding differentially expressed genes (DEGs) specifically identified in the merged dataset by pooling batch 1 data with batch 2 data after MNN correction**. (**a**) Feature enriched GO terms and the corresponding DEGs up-regulated in CMP over GMP. (**b**) Feature enriched GO terms and the corresponding DEGs up-regulated in GMP over CMP. (**c**) Feature enriched GO terms and the corresponding DEGs up-regulated in GMP over MEP. (**d**) Feature enriched GO terms and the corresponding DEGs up-regulated in MEP over GMP.

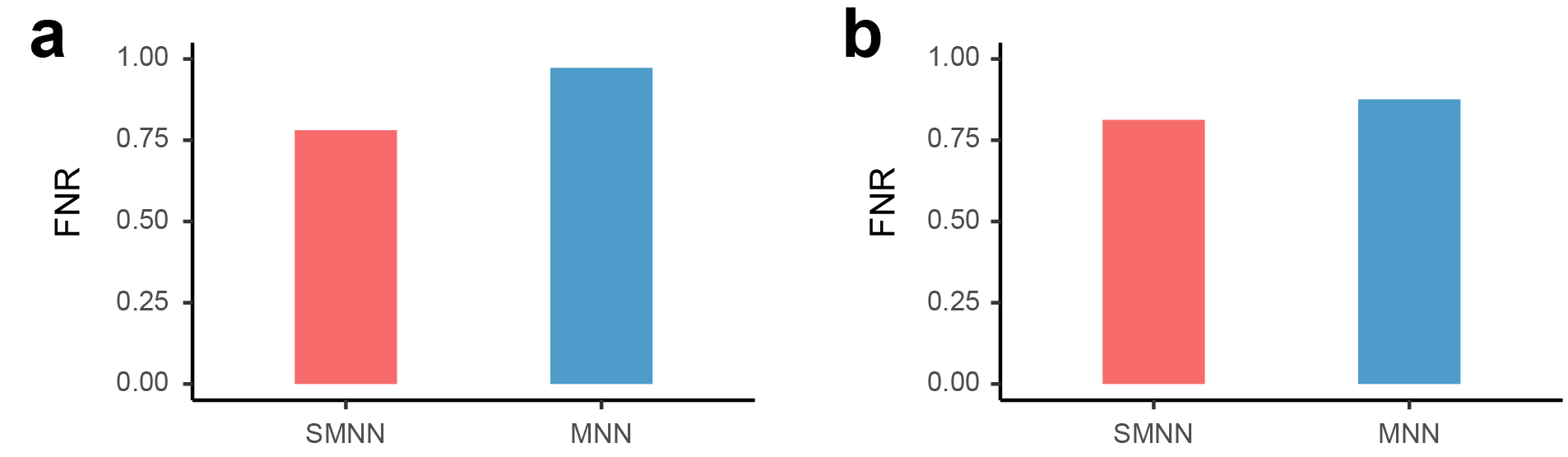

**Fig. S13. False negative rate (FNR) of DEGs (between CMP and GMP), identified in uncorrected batch 1 and in SMNN or MNN-corrected batch 2.** (**a**) FNR of the DEGs up-regulated in CMP over GEP identified in batch 2 after SMNN and MNN correction. (**b**) FNR of the DEGs up-regulated in GMP over CMP identified in batch 2 after SMNN and MNN correction.

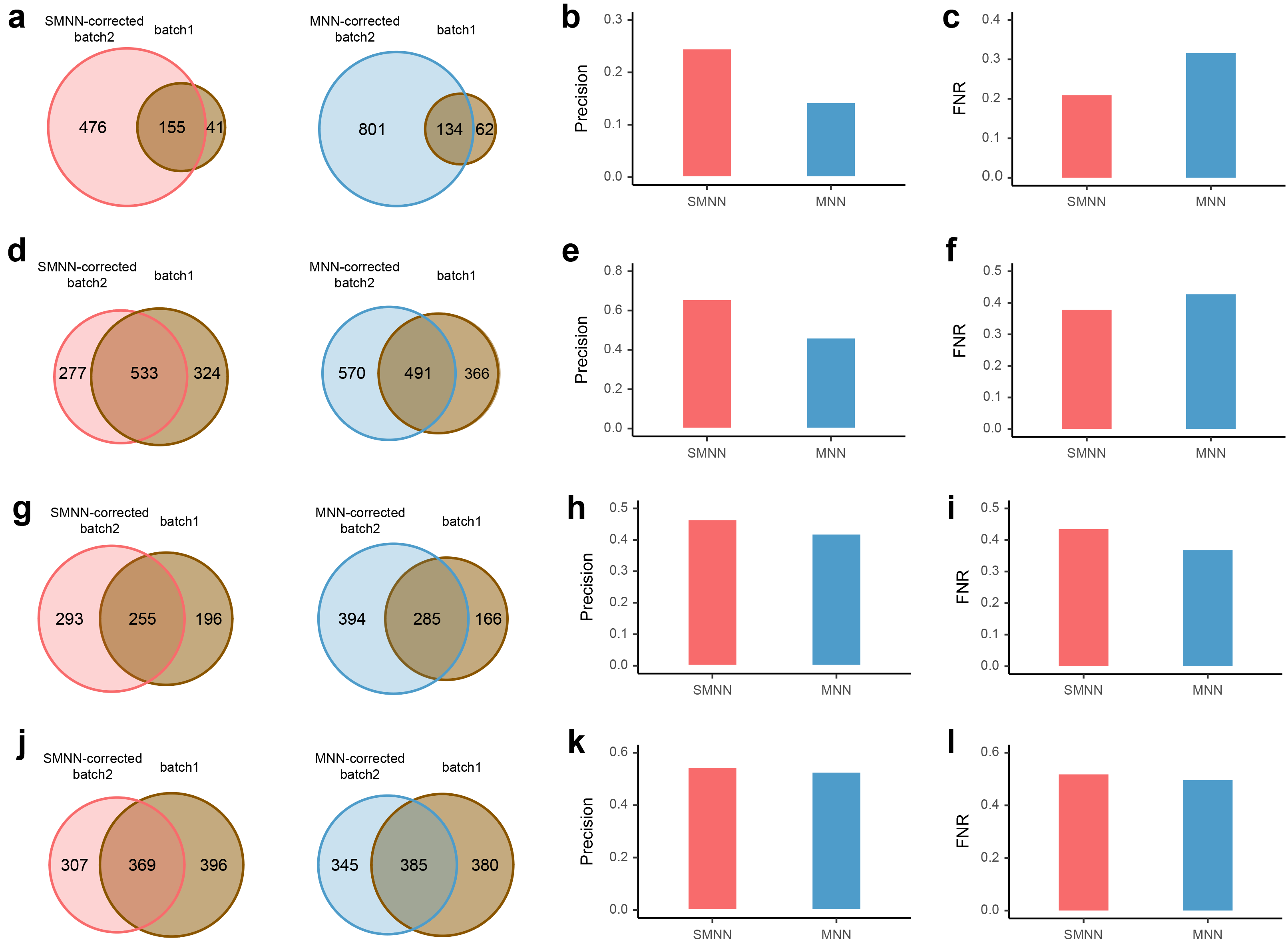

**Fig. S14. Reproducibility of DEGs (between CMP and MEP and between GMP and MEP).** (**a**) Reproducibility of DEGs up-regulated in CMP over MEP. (**b**) and (**c**) Precision and false negative rate (FNR) of the DEGs up-regulated in CMP over MEP. (**d**) Reproducibility of DEGs up-regulated in MEP over CMP. (**e**) and (**f**) Precision and FNR of the DEGs up-regulated in MEP over CMP. (**g**) Reproducibility of DEGs up-regulated in GMP over MEP. (**h**) and (**i**) Precision and FNR of the DEGs up-regulated in GMP over MEP. (**j**) Reproducibility of DEGs up-regulated in MEP over GMP. (**k**) and (**l**) Precision and FNR of the DEGs up-regulated in MEP over GMP.

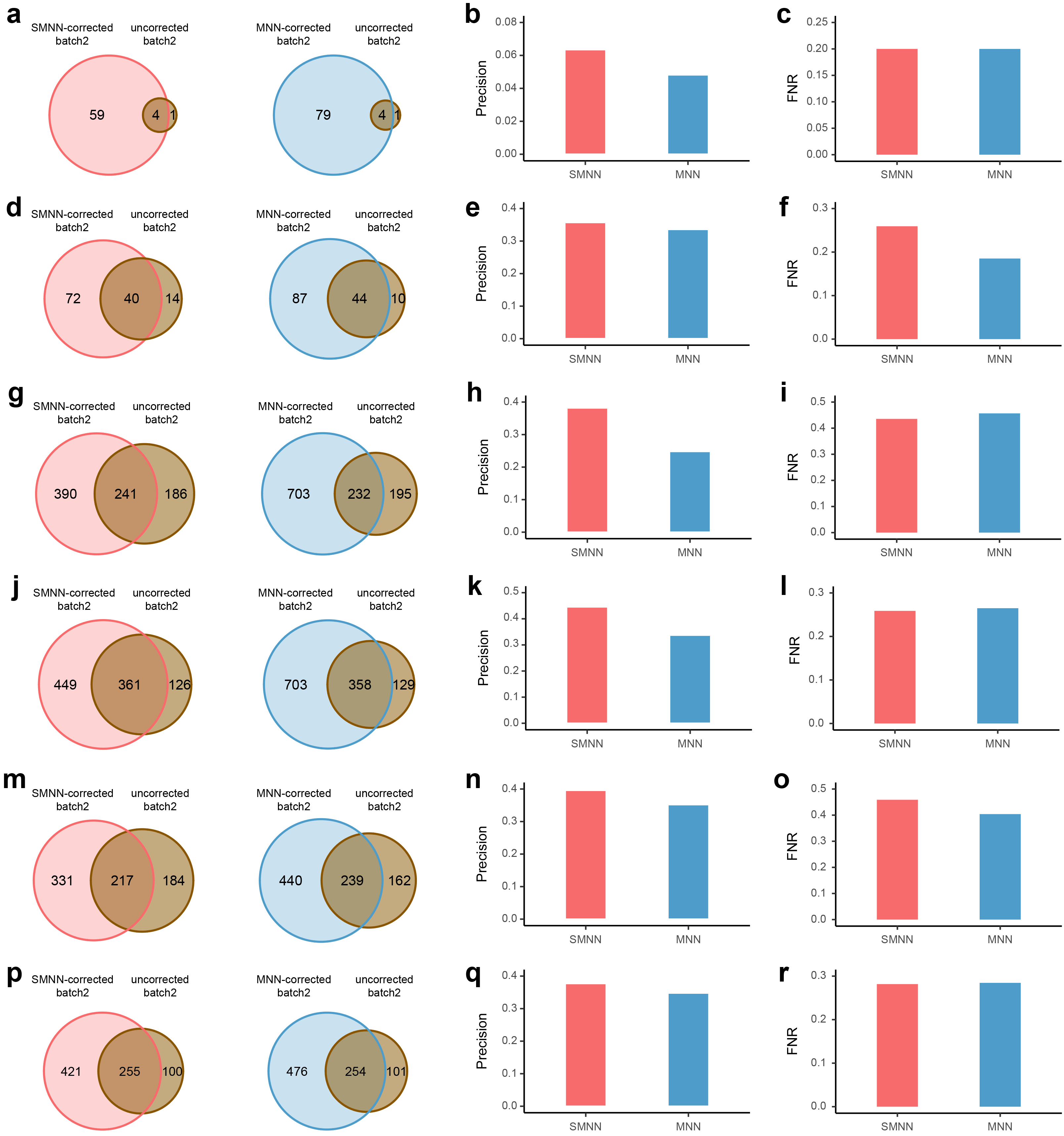

**Fig. S15. Reproducibility of DEGs (between any two cell types, out of the three: CMP, GMP and MEP).** (**a**) reproducibility for DEGs up-regulated in CMP over GMP. (**b**) and (**c**) Precision and false negative rate (FNR) of the DEGs up-regulated in CMP over GMP. (**d**) Reproducibility of DEGs up-regulated in GMP over CMP. (**e**) and (**f**) Precision and FNR of the DEGs up-regulated in GMP over CMP. (**g**) Reproducibility of DEGs up-regulated in CMP over MEP. (**h**) and (**i**) Precision and FNR of the DEGs up-regulated in CMP over MEP. (**j**) Reproducibility of DEGs up-regulated in MEP over CMP. (**k**) and (**l**) Precision and FNR of the DEGs up-regulated in MEP over CMP. (**m**) Reproducibility of DEGs up-regulated in GMP over MEP. (**n**) and (**o**) Precision and FNR of the DEGs up-regulated in GMP over MEP. (**p**) Reproducibility of DEGs up-regulated in MEP over GMP. (**q**) and (**r**) Precision and FNR of the DEGs up-regulated in MEP over GMP.

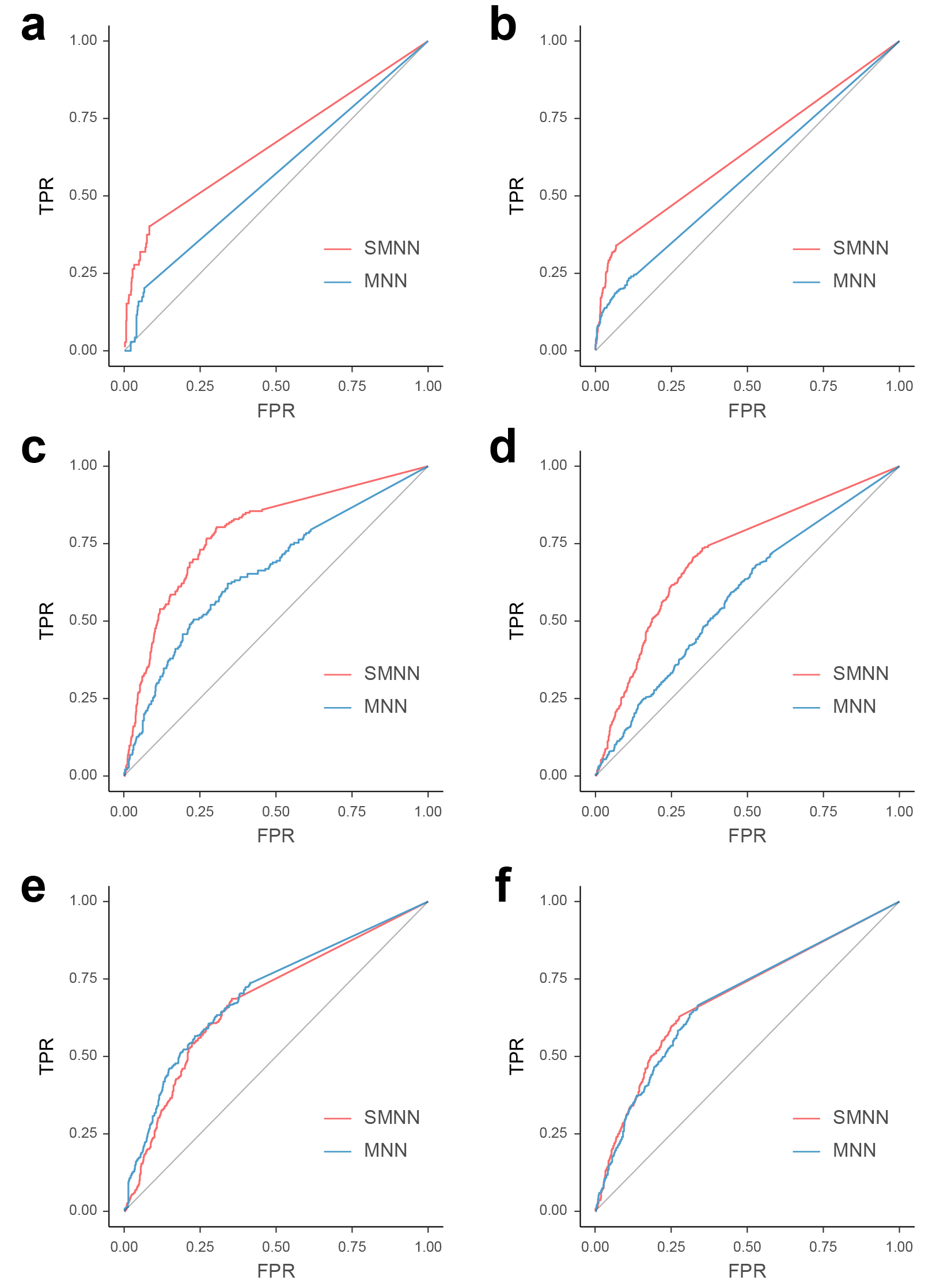

**Fig. S16 ROC curves for DEGs (between any two cell types, out of the three: CMP, GMP and MEP) at various adjusted *p*-value thresholds.** (**a**) ROC curve for DEGs up-regulated in CMP over GMP. (**b**) ROC curve for DEGs up-regulated in GMP over CMP. (**c**) ROC curve for DEGs up-regulated in CMP over MEP. (**d**) ROC curve for DEGs up-regulated in MEP over CMP. (**e**) ROC curve for DEGs up-regulated in GMP over MEP. (**f**) ROC curve for DEGs up-regulated in MEP over GMP.

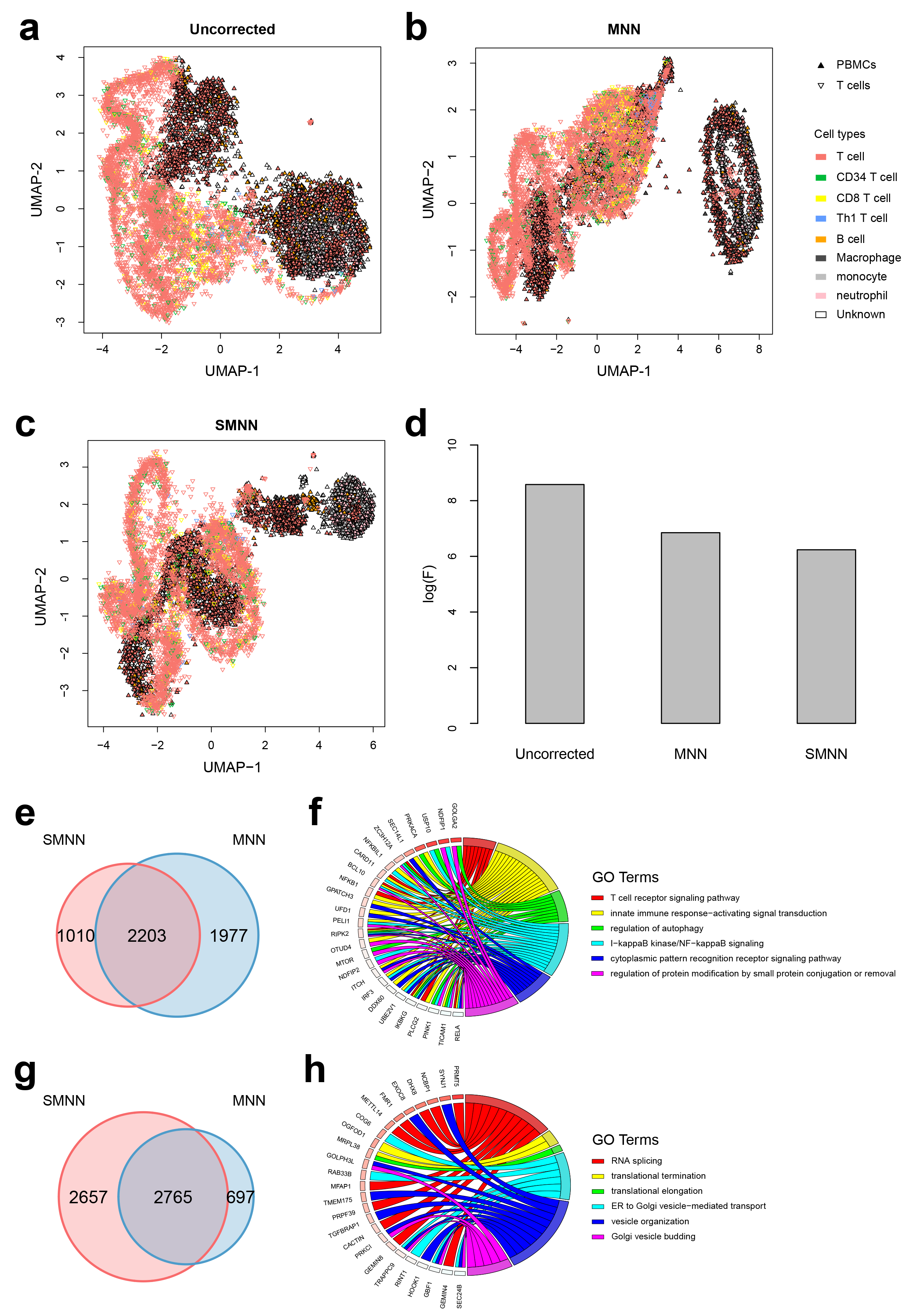

**Fig. S17. Performance comparison between SMNN and MNN in two 10X Genomics datasets for human PBMC and T cells.** (**a-c**) UMAP plots for PBMC and T cells datasets before and after batch effect correction with MNN and SMNN, respectively. Solid and inverted triangle represent PBMC and T cell datasets, respectively; and different cell types are shown in different colors. (**d**) Logarithms of F-statistics for merged data of the two batches. (**e**) Overlap of DEGs up-regulated in T cells over B cells after SMNN and MNN correction. (**f**) Feature enriched GO terms and the corresponding DEGs up-regulated in T cells over B cells. (**g**) Overlap of DEGs up-regulated in B cells over T cells after SMNN and MNN correction. (**h**) Feature enriched GO terms and the corresponding DEGs up-regulated in B cells over T cells.

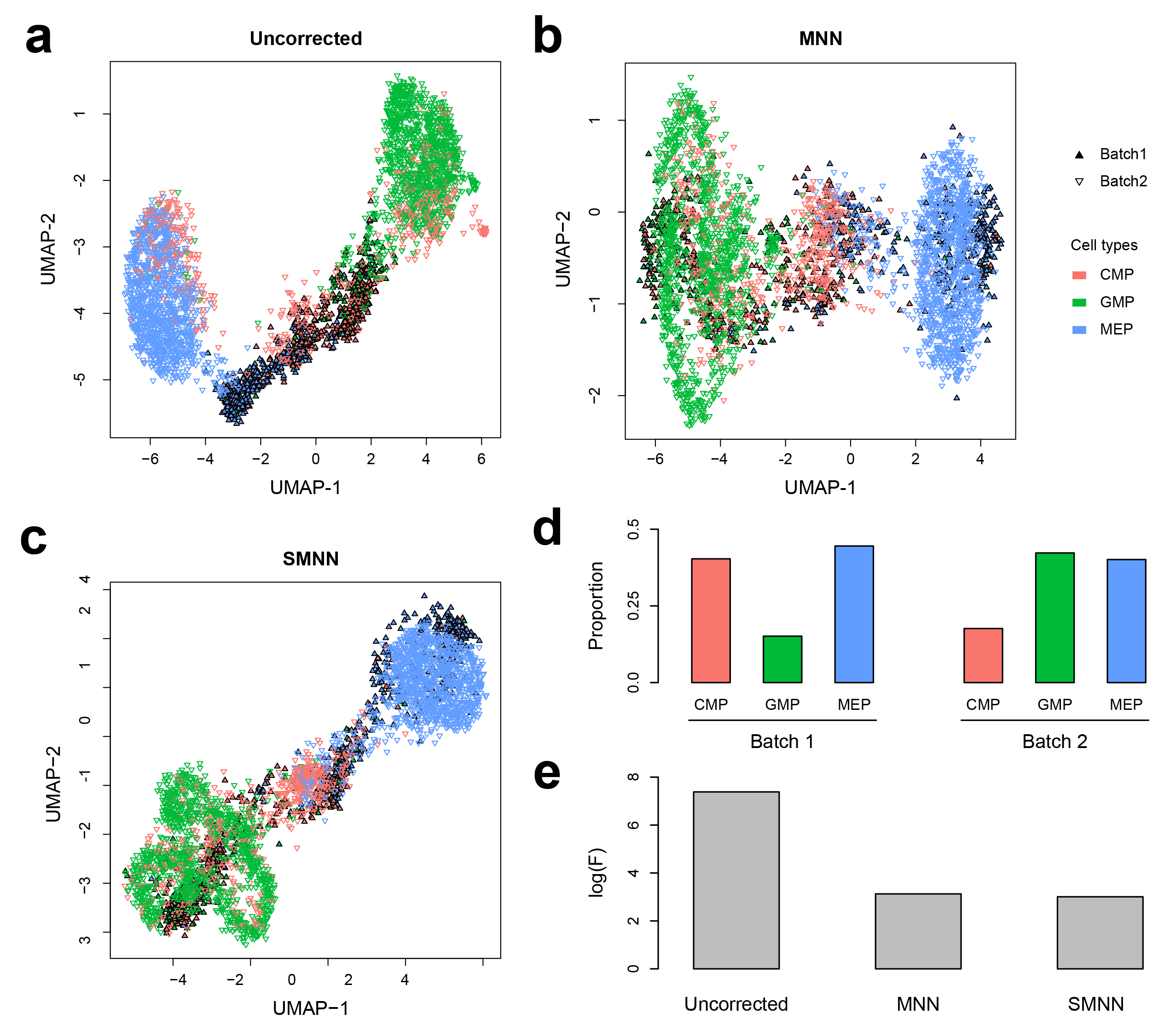

**Fig. S18. Performance of SMNN in two hematopoietic datasets with unbalanced cell population composition.** (**a**-**c**) UMAP plots for two hematopoietic datasets before and after correction with MNN and SMNN, respectively. Solid and inverted triangle represent the first and second batch, respectively; and different cell types are shown in different colors. (**d**) Cell population proportions in the two batches. (**e**) Logarithms of F-statistics for merged data of the two batches before and after correction with MNN and SMNN.
